# Seasonal Patterns of Viromes in Urban Aquatic Environments of Manitoba

**DOI:** 10.1101/2024.03.06.583751

**Authors:** Jhannelle D. Francis, Miguel Uyaguari

## Abstract

Although wastewater and treatment plants harbor many pathogenic organisms’ traditional methods that monitor the microbial quality of wastewater have not changed since the early 1900s and often disregard the presence of other types of significant waterborne pathogens such as viruses. Using advanced technology, our study aims to characterize the taxonomy, functional profiling and seasonal patterns of viral DNA and RNA community structures using metagenomics and quantitative-PCR, for the purpose of establishing the virome distribution in aquatic environment’s receiving wastewater discharge. Environmental water samples were collected at 11 locations in Winnipeg, Manitoba along the Red and Assiniboine Rivers during the Spring, Summer and Fall 2021. Samples were filtered and underwent skimmed milk flocculation for viral concentration.

The taxonomic classification of DNA viruses identified from the RefSeq database (available from MG-RAST) and Kraken 2 Viral Genome database were predominately DNA bacteriophages (*Myoviridae, Podoviridae and Siphoviridae*) which accounted for approximately 90% of each aquatic sample location along the Red and Assiniboine Rivers. Phage related functionalities such as phage tail fiber proteins, phage replication, and phage packaging machinery accounted for 40% of each aquatic samples collected which possibly correspond to the DNA phages that were previously identified. RNA phages such as *Cystoviridae* and *Leviviridae* were identified to a lesser extent accounting for approximately 3 % of each aquatic sample collected, while other viruses such as *Virgaviridae, Retroviridae, Picobirnaviridae* and *Partiviridae* accounted for 7%–100% of aquatic samples. The functionalities of RNA viruses were primarily related to metabolic pathways such as potassium homeostasis, respiratory complexes and sialic acid metabolism, essential for RNA viruses to survive in their host.

**IMPORTANCE:** Municipal wastewater effluents discharged into the Red and Assiniboine Rivers of Winnipeg, Manitoba relies on traditional methods that monitor the microbial quality of wastewater focus solely on the detection of fecal bacteria, which are not necessarily good indicators of viruses or other pathogens. There is also a lack of current wastewater system effluent regulations at the federal and provincial level. Furthermore, previous literature has shown that when viral DNA and RNA sequences are blasted against current genomic databases, approximately 50 % of the viral reads are classified as unknown. The significance of our research in characterizing the virome distribution in aquatic environments addresses a knowledge gap in the current effluent guidelines and a need for regulatory practices. In the long run, fecal indicator bacteria combined with the detection of enteric viruses, may complement assessment of water quality in effluents discharged into rivers.

## INTRODUCTION

Wastewater is defined as contaminated water from a combination of domestic, industrial or agricultural human activities and is typically transported to Wastewater Treatment Plants (WWTPs) for reducing organic load. Wastewater is comprised of liquid and solid waste materials (such as human feces, protein, fat, vegetable and detergents) and is comprised of 99% water and 1% contaminants (Englande et al., 2015). Contaminants of wastewater may generally include approximately 350–1200 mg/L of total solids (TS) waste and 109 number/mL of varying microorganisms such as bacteria, viruses, protozoa, algae, rotifers, nematode and fungi (Warwick et al., 2013).

When municipal wastewater is flushed down the drains from homes and businesses it travels into collection pipes for sanitary sewer systems which transport wastewater to WWTPs for treatment (Environment and Climate Change Canada, 2017). As reported by Statistics Canada in 2017 approximately 5,900 million m^3^ of wastewater is flushed from households, commercial business and industries into WWTPs for treatment where approximately 100 m^3^ of wastewater is processed per day (Statistics Canada, 2019). The City of Winnipeg is located at the intersection of the Red and Assiniboine Rivers and is geographically situated at the center of North America. Wastewater management involves the collection of wastewaters into a city’s sanitary sewer system for transport to WWTPs to reduce organic matter. After which, local sewer pipelines in Winnipeg direct the wastewater to one of three WWTPs for treatment.

The city of Winnipeg with a population of 841,000 (as of September 2023) has three major WWTPs: North End Water Pollution Control Centre (NEWPCC), South End Water Pollution Control Centre (SEWPCC) and West End Water Pollution Control Center (WEWPCC) (City of Winnipeg, 2021). All three sewage plants combined discharge a total of 270 million liters of effluents per day into local rivers. The three primary stages of wastewater treatment include: primary/physical, secondary/biological, and tertiary/disinfection treatment using UV light (City of Winnipeg, 2021).

Fecal indicator bacteria are the current gold standard of aquatic health when assessing the microbial quality of wastewater, an approach that has been used for over a century. Fecal coliform bacteria are microscopic organisms that originate in the intestines of warm-blooded animals and can also be present in fecal material excreted from the intestinal tract (Khan, F. M., 2020, Rodrigues et al., 2017). Domestic wastewater typically contains a higher number of coliform bacteria from fecal origin (*Escherichia coli*) than those from non-fecal origin due to the accumulation of fecal material of humans or other animals (Gokul et al., 2019). However fecal coliform bacteria can enter surface waters primarily through anthropogenic activities by direct discharge of human sewage from WWTPs, agricultural practices or from storm runoffs that transport animal wastes to streams through storm sewers. Treatments designed to screen and remove fecal coliform, *E. coli* and waterborne parasites *Cryptosporidium* and *Giardia* from wastewater at Winnipeg’s WWTPs may include chlorine with UV light disinfection. In compliance to ensure the removal of fecal bacteria and coliforms from wastewater with the province of Manitoba’s recreational guidelines, UV radiation at ∼254 nm lasts for 4 seconds on average and can last for 2 seconds during peak flow (City of Winnipeg, 2021).

The presence of fecal coliforms in effluents represents inefficient removal of fecal material that may promote growth of organic matter and deplete the oxygen available to aquatic species in local rivers (Environmental Protection Agency, 2012). While *E. coli* has been widely-adopted across many treatment plants as an indicator of fecal contamination for water quality, it limits the amount of information derived from a negative-culture result. When a treated wastewater sample is tested for the presence of fecal indicator bacteria, a negative culture result does not rule out the presence of other significant types of waterborne pathogens such as viruses or protozoans (Ramírez-Castillo et al., 2015). Whereas a positive culture result test may be indicative of recent fecal contamination it does not provide information on the source of contamination or on the risk to health. The presence of contaminants and pathogenic microorganisms can pose detrimental health risks to humans who rely on surface waters for domestic and recreational use. Depending on the type of pathogen, chemicals and other toxic substances present, the severity of diseases incurred will vary. Many reported cases of enteric diseases include gastrointestinal related illnesses such as vomiting, nausea, fever, abdominal pain and loss of appetite all of which are relatively mild with a short recovery time (Government of Canada, 2017). Other acute illnesses may include meningitis, poliomyelitis and non-specific febrile illnesses (Government of Canada, 2013).

Studies suggest that there has been an increasing rise in the amount of pollutants present in effluent samples compared to untreated wastewater, possibly due to the lack of sufficient treatment processes at WWTPs (Rodrigues et al., 2017, Zhang et al., 2009; Wear et al., 2021). Without receiving adequate treatment at WWTPs, wastewater pollutants such as antibiotics and their metabolites, bacteria, viruses, protozoa, heavy metals, and hormones can adversely affect the aquatic environment to which it is discharged into. Between 1998 and 2003, 18 norovirus outbreaks occurred in Finland impacting more than 10,000 residents due to inadequate wastewater treatment and further contamination of groundwaters or surface waters (Maunula et al., 2005). Even though treated water samples tested negative for the presence of *E. coli* and total coliforms, these outbreaks raised questions about the sole usefulness of the current microbial standards of aquatic health for routine wastewater quality testing. Aside from Norovirus outbreaks, there are many other reports on Rotavirus, crAssphage, and Adenovirus prevalence in freshwaters (Hellmer et al., 2014; Garcia et al., 2022; Mafumo et al., 2023).

When compared to *E. coli* and coliforms there is a reduction efficiency of viruses at WWTPs (Ito et al., 2016; Tandukar et al., 2021). Moreover, there is also a lack of current wastewater system effluent regulations at the federal and provincial level. Instead, it is only advised that effluents should contain less than 200 coliform units per 100 mL of water (Justice Laws, 2015). Therefore, this study aimed to address a knowledge gap in the current effluent guidelines and a need for regulatory practices. Waterborne enteric viral outbreaks occurred in groundwaters of Alberta and Ontario at 11 % positive sample frequency in 2013 (Government of Canada, 2017d). However, some viral outbreaks go unperceived or not reported. While there are cases of viral outbreaks, viruses in aquatic ecosystems are not studied in detail due to a lack of known viral databases. Previous literature has shown that when viral DNA and RNA sequences are blasted against viral genome databases, approximately 50 % of the viral reads are classified as unknown (Goodacre et al., 2018; Tisza and Buck, 2021). Since viral DNA and RNA sequences in current genomic databases are classified as unknown this represents an opportunity for this thesis to identify viral fractions for the purpose of establishing the virome distribution in aquatic environment’s receiving wastewater discharge.

There is a need for additional water and wastewater surveillance and detection of microorganisms that this work intends to leverage on. A better understanding of temporal changes in changes of viromes (including enteric viruses) in surface waters is equally as important as fecal indicator bacteria, in assessing the utility of current biomarkers of fecal contamination. During the pandemic wastewater surveillance emerged as a monitoring tool to screen for SARS-CoV-2. Surveillance of this sentinel system is strongly suggested to be expanded to other pathogens as public health preparedness. Canada and its public health agencies should expand their surveillance capability. Results from the proposed study will help to be prepared for future threats to anticipate and prevent the spread of antimicrobial diseases. Our research serves as an exploratory study with the primary objective to characterize the DNA and RNA virome distribution in urban influenced environments using metagenomics.

## RESULTS

### High-throughput screening revealed that urban aquatic environments of Manitoba contained predominately DNA bacteriophages with phage-related functionalities

Taxonomic classifications of assembled reads identified from NCBI RefSeq database (available at MG-RAST; Accession Number: PRJNA1011997) at the family level revealed an abundance of DNA viruses such as *Myoviridae, Podoviridae* and *Siphoviridae* which accounted for the following relative abundances within aquatic samples collected during the Spring, Summer and Fall of 2021: 15%–30%, 20%–50% and 20%–60% respectively of (Fig. 1A). Although these viruses were also identified in Kraken 2 viral genome database, their abundance differed and accounted for approximately 30%–50%, 9%–17% and 30%–55% respectively of the aquatic samples collected (Fig. 1B). Other viruses such as *Herpesviridae* and *Baculoviridae* DNA viruses were abundant to a lesser extent with each representing approximately 3 % of the aquatic samples collected (Fig. 1B). Assembled reads of DNA viruses were uploaded into MG-RAST SEED Subsystems database and revealed highest abundances (%) of the following level 3 functionalities: phage tail fiber proteins (5%– 40%), phage replication (55%–38%), phage packaging machinery (5%–33%) and phage entry and exit functionality (6%–17%), in aquatic samples 1-11 collected along the Red and Assiniboine Rivers (Fig. 2). Whereas these functionalities were abundant in all seasons, other functionalities such as folate biosynthesis (6%–13%) and gene transfer agent (6%–27%) were most abundant in the Spring and Summer only.

**FIG 1:**
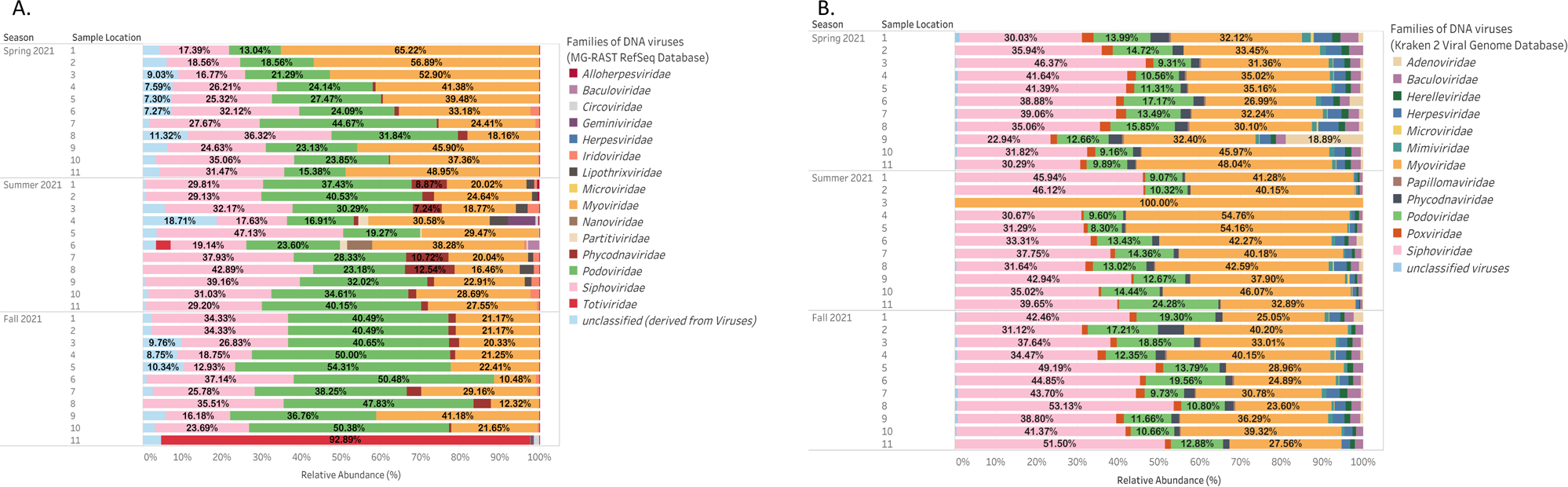
Observational trends depicting taxonomic families of DNA viruses during the Spring Summer and Fall of 2021 that were identified from (A) NCBI RefSeq database (available at MG-RAST) and (B) Kraken 2 Classifier Viral Genome Database.

**FIG 2:**
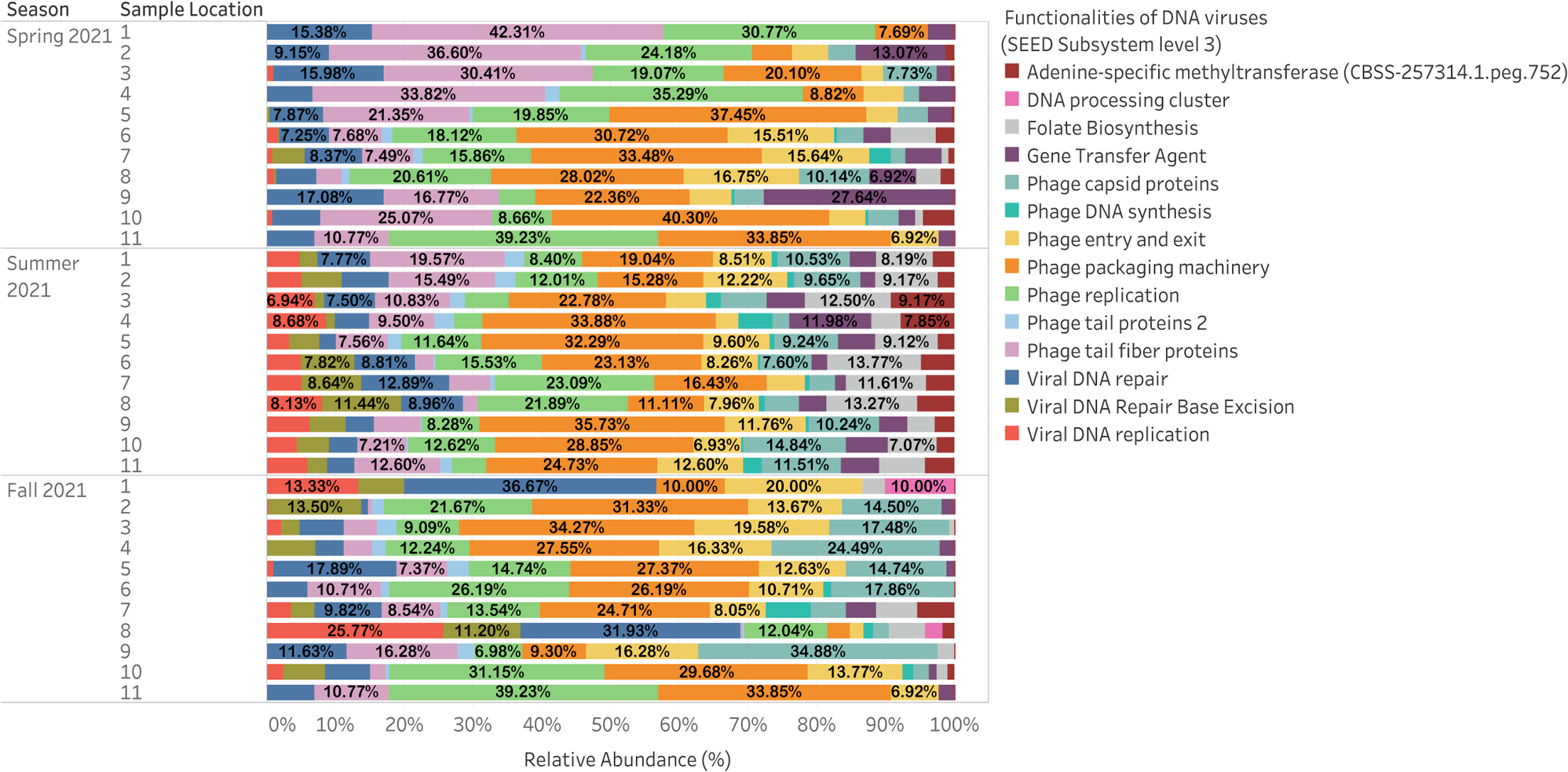
Observational trends depicting level 3 Functionalities for DNA viruses during the Spring, Summer and Fall of 2021. Functionalities were identified from MG-RAST SEED Subsystem database (level 3).

### RNA viruses present in urban aquatic environments of Manitoba fluctuated across seasons with a large proportion remaining unclassified and evidenced functionalities primarily related to metabolic pathways

RNA viruses identified from NCBI RefSeq database (available at MG-RAST, Accession Number: PRJNA1011997) at the family level revealed an abundance (%) of *Retroviridae* (7%–100%) and *Tombusviridae* (13%–75%) in Spring 2021, *Partiviridae* (12%–80%) and *Picobirnaviridae* (7%–75%) in summer 2021 and *Reoviridae* (9%– 38%) and *Picobirnaviridae* (8%-32%) in Fall 2021 across all aquatic samples collected (Fig. 3A). RNA viruses identified from Kraken 2 viral genome database revealed an abundance (%) of *Retroviridae* (36%–97%) in Spring 2021, *Virgaviridae* (64%–88%), *Retroviridae* (8%–97 %) and *Partiviridae* (7%–75%) in Summer 2021 and *Virgaviridae* (5%–71%), *Retroviridae* (9%–65%) and *Picobirnaviridae* (9%–26%) in Fall 2021 across all surface water samples collected (Fig. 3B). The findings from NCBI RefSeq database (available at MG-RAST; Accession Number: PRJNA1011997) and Kraken 2 viral genome database suggest that the DNA viruses identified were relatively more stable and consistent throughout changes in seasonal weather whereas the RNA viruses identified displayed a greater degree of seasonal variability and disparity. Unclassified DNA and RNA viromes were present in all aquatic samples with a higher percentage of viral RNA remaining unclassified than that of DNA. A higher percentage of viral genomes remained unclassified in NCBI RefSeq database (available at MG-RAST; Accession Number: PRJNA1011997) (4%–100%) compared to that of Kraken viral genome database (11%–50%) across all aquatic samples collected. Level 3 functionalities identified from assembled reads of RNA viruses revealed a consistent abundance (%) of respiratory complexes (6%–44%), potassium homeostasis (4%–25%), sialic acid metabolism (5%–25%) and purine conversions (5%–50%), in each season from the aquatic samples collected 1-11 along the Red and Assiniboine Rivers (Fig 4). In Spring 2021 other functionalities such as tRNA pseudouridine synthase B (6%–30%), fatty acid biosynthesis FASII (4%–11%), methionine biosynthesis (6%–13%), HtrA and Sec secretions (9%–17%) and ABC transporters (5%–16%) were also found to be abundant. Cytochrome c oxidase (36%–52%) was found to be abundant in the Summer 2021. Other functionalities such as tRNA pseudouridine synthase B (6%– 60%), triacylglycerol metabolism (14%–41%) and tRNA processing (4%–14%) were also found to be abundant in Fall of 2021. Whereas the level 3 functionalities of DNA viruses appeared to be stable throughout each season, those of RNA viruses were found to be seasonally variable.

**FIG 3:**
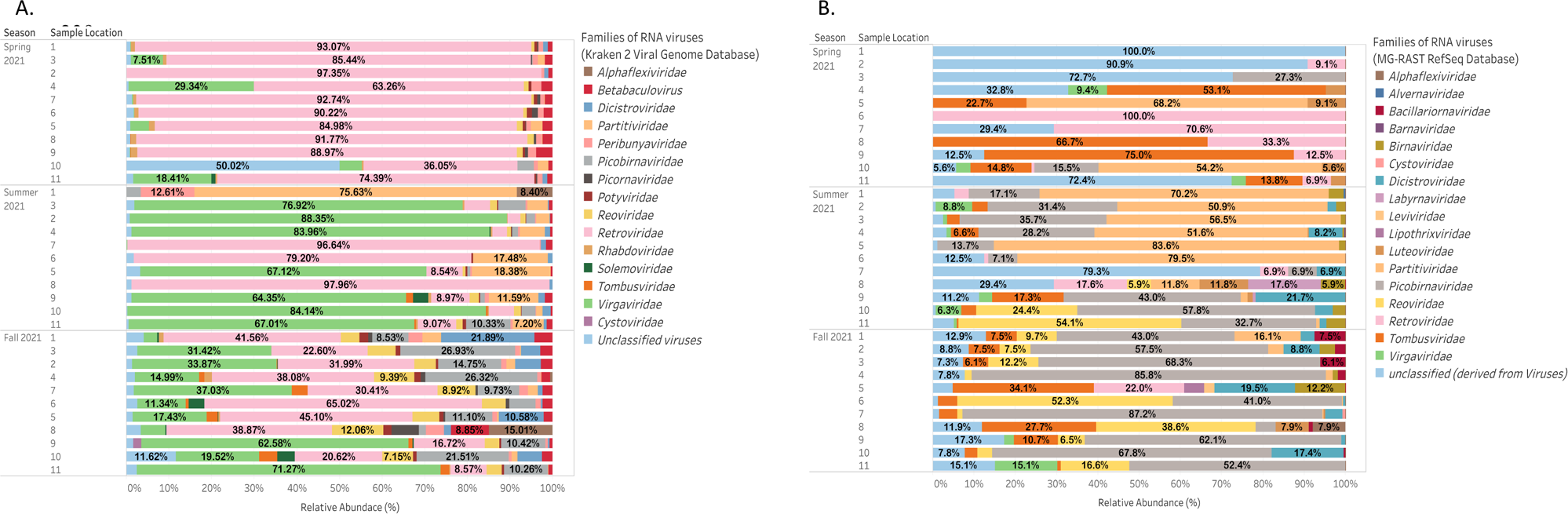
Observational trends depicting taxonomic families of RNA viruses during the Spring, Summer and Fall of 2021 that were identified from (A) NCBI RefSeq database (available at MG-RAST) and (B) Kraken 2 Classifier Viral Genome Database

**FIG 4:**
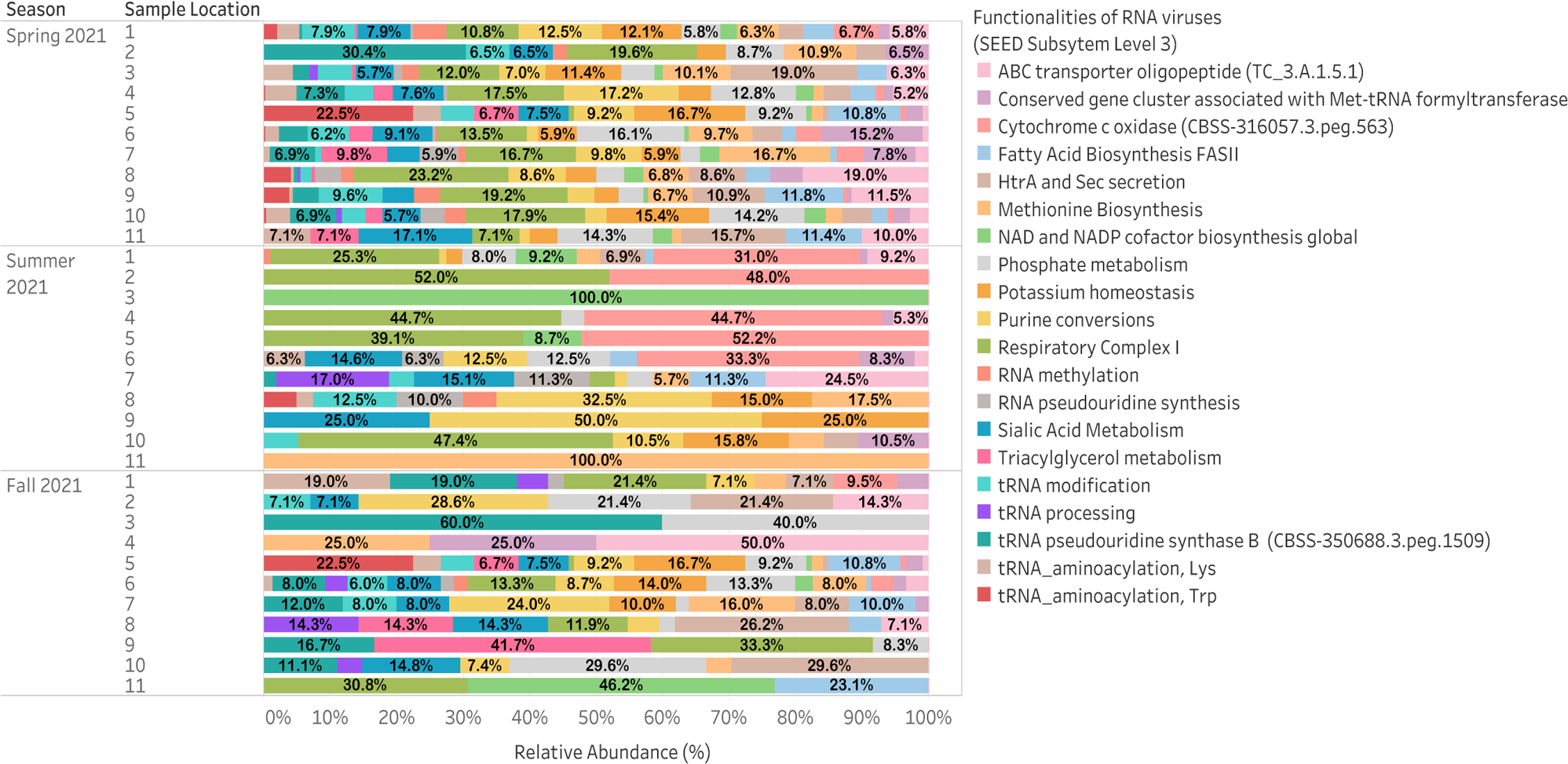
Observational trends depicting level 3 Functionalities for RNA viruses during the Spring, Summer and Fall of 2021. Functionalities were identified from MG-RAST SEED Subsystem database (level 3)

### Exploratory factor analysis (orthomax rotation) revealed that daylength, temperature and land use from anthropogenetic activities affected were significant factors affecting relative abundances of urban viral communities in Manitoba

Factor 1 or “land use” accounted for 19.53 % of the observed variability, while that factor 2 or “daylength and temperature” accounted for 16.17 % of the observed variability. Three distinct clusters of sites 6, 7 and 8 from the Assiniboine River are each visually identified for the Spring, Summer and Fall of 2021 collection events (Fig. 5). The remaining sample locations 1-5 and 9-11 from the Red River appeared to form individual clusters for the Summer and Fall 2021 collection events each while those for the Fall collection event was scattered. Factor loadings for the environmental parameters – total phosphorous (TP), five-day biochemical oxygen demand (BOD_5_), TN, river water flow rate and phosphate (PO_4_) all fell within the same cluster as sites 1-5 and 9-11 for the Fall 2021 sample collection. On the other hand, rainfall and carbonaceous biochemical oxygen demand (cBOD_5_) were within the same quadrant as sites 1-5 and 9-11 from the Summer collection event. HAdV and crAssphage occurrence in aquatic environments were both evidenced to be significantly impacted and positively correlated to the following water quality parameters: temperature (r = 0.567, p = 6.00E-04; r = 0.452, p = 8.3E-03), BOD_5_ (r = 0.521, p = 1.90E-03; r = 0.627, p <1.00E-04), daylength (r = 0.790, p <1.00E-04; r = 0.421, p = 1.46E-02) and *E. coli* counts (r = 0.615, p <1.00E-04; r = 0.346, p = 4.83E-02), while also negatively impacted by precipitation (r = −0.735, p <1.00E-04; r = - 0.295, p = 6.56E-02) (Fig. 6). crAssphage was also found to be significantly impacted and positively correlated to NH4-N and TP (r = 0.492, p = 3.60E-02 and r = 0.509, p = 2.4E-02 respectively).

**FIG 5:**
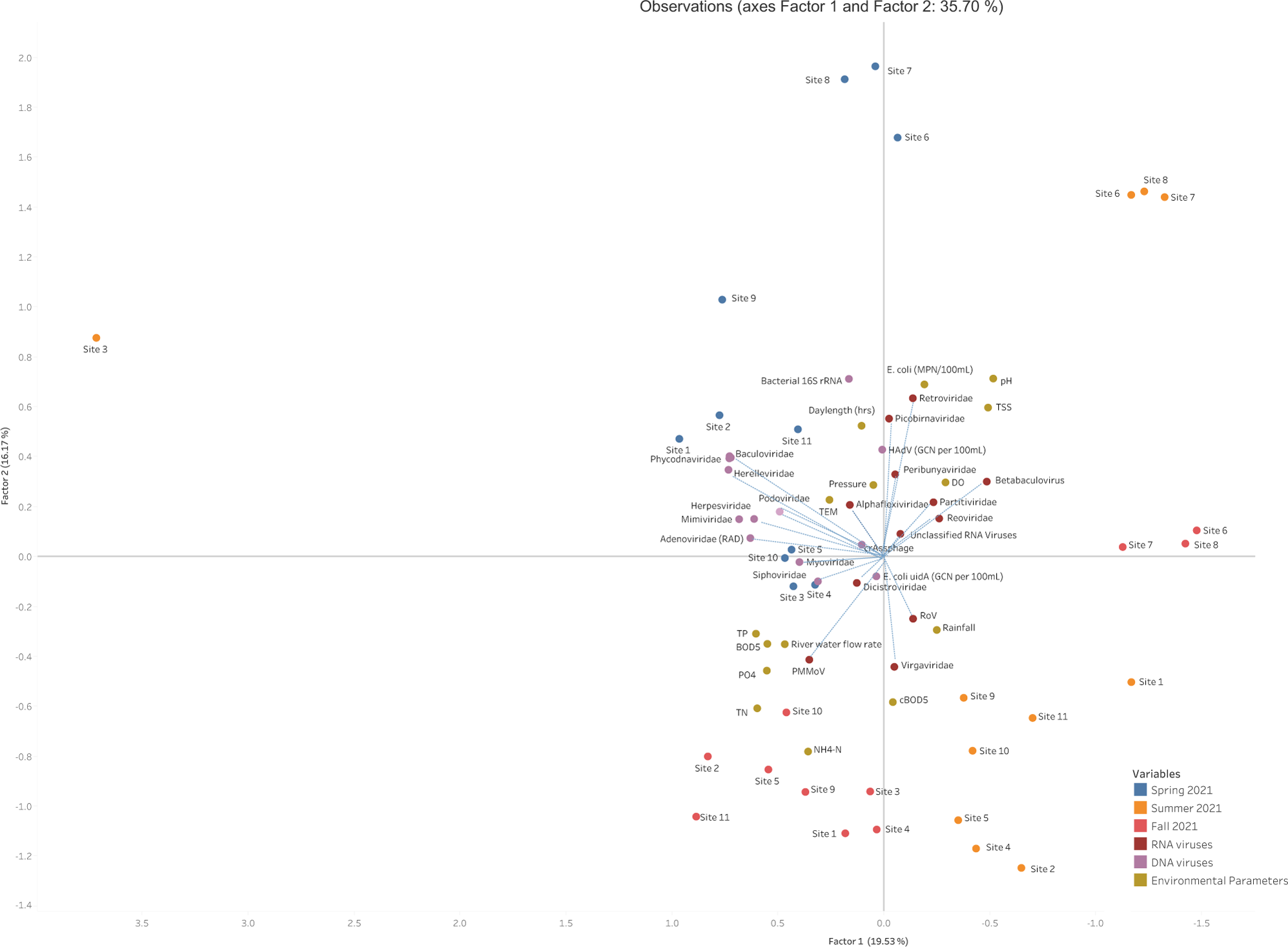
Factor analysis of viral DNA and RNA families and environmental variables observed at each sample collection site (1-11) during the Spring (blue dots), Summer (orange dots) and Fall (red dots) of 2021. Factor 1 represents land use which incorporates urban influenced watersheds on DNA viruses and agricultural influenced watershed on RNA viruses while factor 2 represents the water quality parameters daylength and temperature. TSS, total suspension solids; DO, dissolved oxygen; TEM, temperature; cBOD_5_, carbonaceous biochemical oxygen demand; BOD, biochemical oxygen demand; TP, total phosphorous; TN, total nitrogen; NH4-N, ammonium, PO_4_, phosphorous; *E. coli* (MPN/100 mL), *E. coli* colony forming unit counts. Blue dashed lines represent factor loading values for viral families.

**FIG 6:**
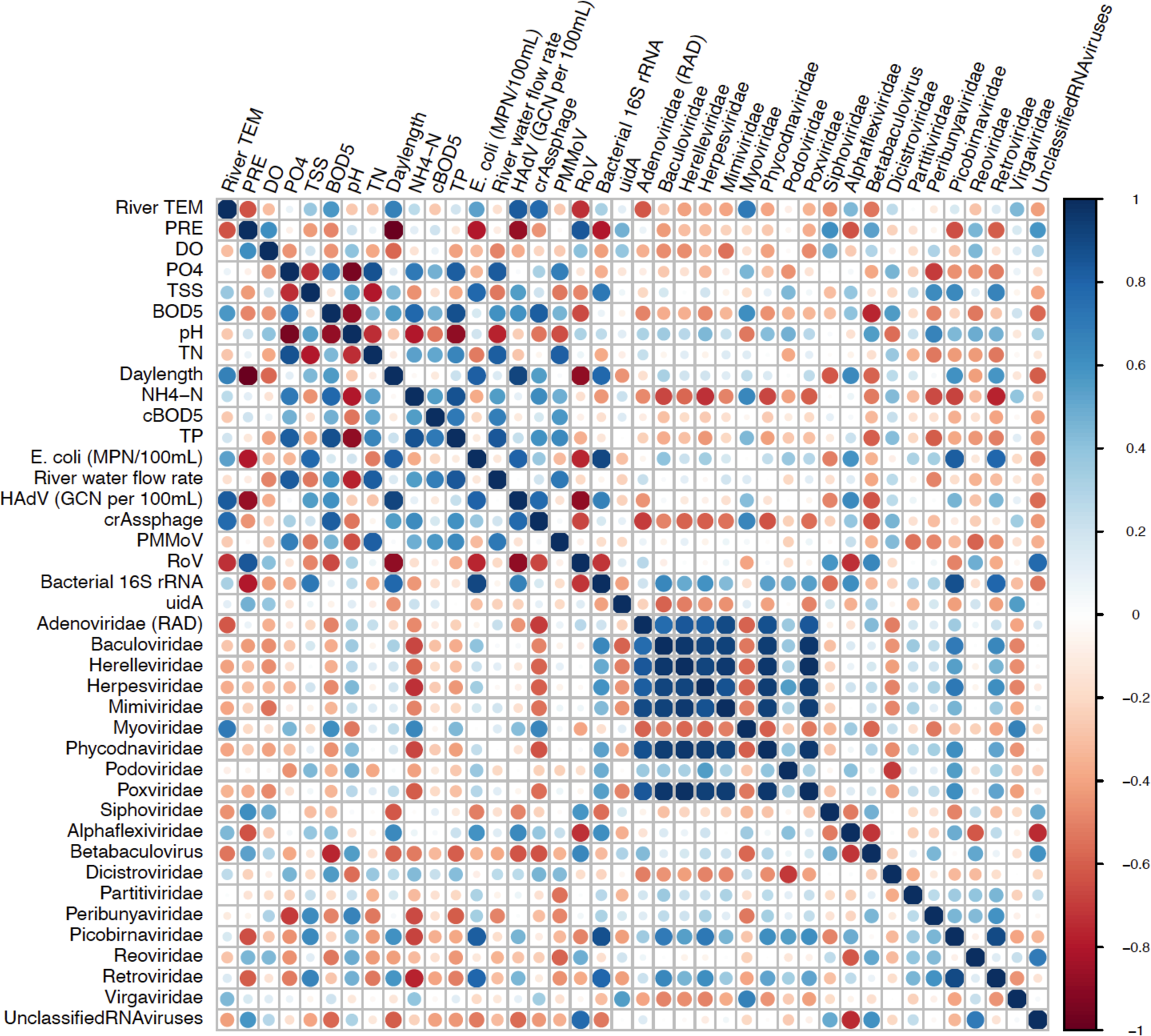
Correlogram of viral DNA and RNA families and environmental variables observed at each sample collection site (1-11). Correlation coefficients range from +1 to −1 and are represented by circles colored blue to red. TSS, total suspension solids; DO, dissolved oxygen; PRE, precipitation; TEM, temperature; cBOD_5_, carbonaceous biochemical oxygen demand; BOD, biochemical oxygen demand; TP, total phosphorous; TN, total nitrogen; NH4-N, ammonia, PO_4_, phosphorous; *E. coli* (MPN/100mL), *E. coli* colony forming unit counts. Adenoviridae RAD was identified metagenomically while HAdV GCN per 100 mL was assessed though quantitative analyses.

Whereas the abundance of DNA enteric viruses was both affected by similar water quality parameters, RNA enteric viruses such as PMMoV and RoV were influenced by different parameters. In Fig. 7, the presence of PMMoV in surface waters was found to be significantly impacted and positively correlated to PO_4_ (r = 0.361, p = 3.92E-02), TN (r = 0.516, p = 0.210), TP (r = 0.347, p = 4.78E-02). In contrast, the abundance of RoV was evidenced to be negatively influenced by temperature (r = - 0.393, p = 2.36E-02), TSS (r = −0.344, p = 4.98E-02), BOD_5_ (r = −0.495, p = 3.40E-03), daylength (r = −0.650, p = 1.00E-04) and *E. coli* counts (r = −0.457, p = 7.50E-03) and positively correlated to precipitation (r = 0.593, p = 3.00E-04) (Fig. 6). Aside from PMMoV, the concentration of PO_4_ in surface waters was also revealed to have a significant negative correlation with *Peribunyaviridae* (r = −0.385, p = 2.67E-02). The temperature of surface waters was also found to positively affect the abundance of other viruses such as *Myoviridae* (r = 0.536, p = 1.30E-03) and *Virgaviridae* (r = 0.344, p = 4.99E-02) while precipitation was also found to negatively impact the abundance of *Herelleviridae* (r = −0.308, p = 8.10E-02), *Picobirnaviridae* (r = −0.407, p = 1.86E-02) and *Retroviridae* (−0.4.43, p = 9.90E-03). *Baculoviridae* was found to be the only virus positively (r = 0.344, p = 4.90E-02) affected by temperature and negatively affected by NH4-N (r = −0.364, p = 3.68E-02). The presence of *Picobirnaviridae* and *Retroviridae* were also both positively impacted by daylength (r = 0.451, p = 8.30E-03; r = 0.497, p = 3.30E-03) and *E. coli* counts (r = 0.609, p = 2.00E-04; r = 0.549, p = 9.00E-04) respectively. Aside from *Baculoviridae,* the concentration of NH4-N in aquatic environments revealed to have a significant and positive impact on other viruses such as *Herpesviridae* (r = 0.383, p = 9.60E-03), *Myoviridae* (r = 0.383, p = 2.79E-02) and *Poxviridae* (r = −0.387, p = 2.62E-02) with also a strong negative correlation to *Retroviridae* (r = −0.606, p = 2.00E-04) (Fig. 6).

**FIG 7:**
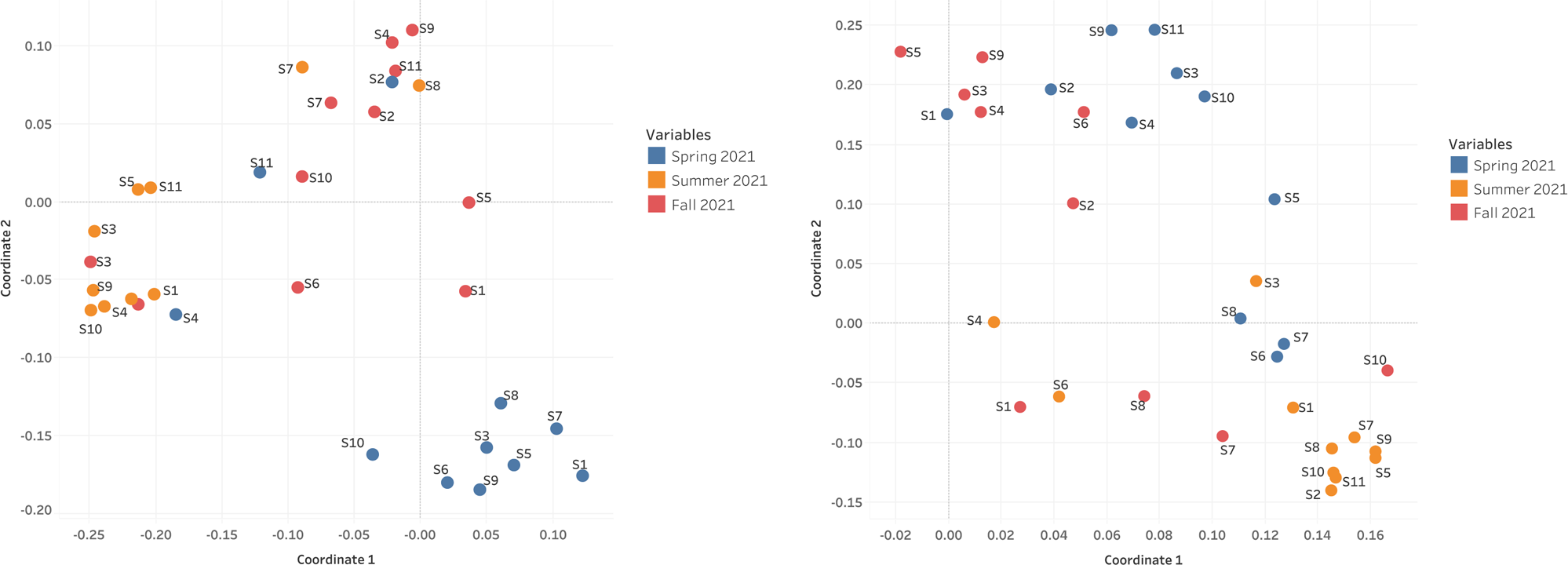
Assessment of the beta diversity among (A) DNA and (B) RNA viral communities present in aquatic samples 1-11 collected during the spring (blue dots), Summer (orange dots) and Fall (red dots) 2021. The Bray-Curtis dissimilarity (beta diversity) index were calculated from the outputs of the taxonomic classifications generated by MG-RAST server.

Whereas the concentration of BOD_5_ was negatively correlated with *Betabaculovirus* (r = −0.386, p = 2.64E-02), TSS was found to be positively correlated with *Peribunyaviridae* (r = 0.452, p = 8.20E-03), *Retroviridae* (r = 0.366, p = 3.63E-02) and *Picobirnaviridae* (r = 0.358, p = 4.09E-02). Interestingly, pH did not appear to have a major influence on the abundance of viruses in the Red and Assiniboine Rivers, while DO appear to have the least impact on the viral community as *Mimiviridae* was the only virus whose abundance was negatively affected (r = −0.447, p = 9.10E-03) (Fig. 5; Fig. 6). Other parameters such as river water flow rate and TP revealed to only have a significant impact on the enteric viruses that were quantitatively assessed in the study, while cBOD_5_ was evidenced to be an insignificant parameter on the abundances of all viruses assessed within the study (Fig. 5; Fig. 6). Expectedly, bacterial *16S rRNA* was significantly positively correlated to *E. coli* counts (MPN/100 mL) (r = 0.690, p <1.00E-04) (Fig. 6). However, *E. coli* counts (MPN/100 mL) was also found to have a surprisingly significant impact and positive correlation to *Picobirnaviridae* (r = 0.609, p = 2.00E-04) (Fig. 6). A strong positive correlation was found between bacteria and *Baculoviridae* (r = 0.449, p = 8.70E-03), *Herpesviridae* (r = 0.441, p = 1.03E-02), *Phycodnaviridae* (r = 0.464, p = 6.50E-03), Herelleviridae (0.372, p = 3.30E-02), *Alphaflexiviridae* (r = 0.392, p = 2.41E-02), *Retroviridae* (r =0.539, p = 1.20E-03), *Picobirnaviridae* (r = 0.560, p = 7.00E-04) and HAdV (r = 0.540; p = 1.20E-03) (Fig. 5; Fig. 6). Bacterial counts (assessed via qPCR) were found to negatively impact the abundance of RoV (r = −0.543, p = 1.10E-03) (Fig. 5; Fig. 6). It is also worthy to note that the unclassified RNA viruses did not appear to be significantly impacted by any of the water quality parameters included in the study.

### Beta diversity of DNA and RNA virome distribution were influenced by seasonality

The populations of DNA viruses present during the Spring and Fall were evidenced to be more closely related (low Bray-Curtis dissimilarity distance of 2.71E-01 compared to those present in the Summer of 2021 (Fig. 7A). DNA viruses observed during the Summer appeared to be more diverse from those identified during the Spring and Fall with high Bray-Curtis dissimilarity distances of 1.84 and 2.21 respectively (Fig. 7A). In contrast, the RNA viral communities identified in the Summer, appeared to be most similar and closely related (a low Bray-Curtis dissimilarity distance of 4.08E-02) to those identified in the Fall of 2021 (Fig. 7B). The RNA viruses identified in the Spring of 2021 were the most diverse from those identified in the Summer (Bray-Curtis dissimilarity distance of −1.17). These viruses present during the Spring were also observed to be dissimilar from those identified during the Fall (high Bray-Curtis dissimilarity distance of −1.06).

### Alpha diversities Simpson’s (1-D), Shannon (H) and Chao-1 reveal seasonal patterns amongst DNA and RNA viral communities in urban Manitoba

Using the Simpson’s (1-D) diversity index, the DNA viral communities present in each surface water sample collected at eleven locations (1-11) along the Red and Assiniboine River evidenced more species diversity during the Spring and Summer of 2021, while samples collected in the south-end (location numbered 2) and in the north end (location numbered 11) of the Red River collected in Fall of 2021 were less diverse (Table 1; Table 2; Fig. 8). To further confirm the species diversity and richness observed in the sequenced samples, the Shannon diversity index was measured. Similarly, the DNA viruses present in each surface water samples collected during the Spring and Fall measured high values of Shannon diversity index which indicated high species diversity present in these samples (Table 1). However, in Fall of 2021 DNA viral communities amongst the samples collected measured lower values of Shannon diversity index. Although the results of a Kruskal-Wallis Test for Shannon and Simpson diversity indices revealed that there were no statistical differences (p–value ≥ 0.05) in the diversity of species of DNA viruses between sample locations, statistical differences (p-value ≤ 1.00E-04) in specie diversities of DNA viruses were observed between the seasons Spring, Summer and Fall of 2021 when the samples were collected. To assess the abundance of viruses present in each sample (1-11) collected, Chao-1 richness index was measured. Samples 6 and 7 collected at the Assiniboine River in the spring and fall respectively, and sample 2 collected at the south end of the Red River in Summer of 2021, all evidenced DNA viruses that were most abundant in comparison to those identified from the remaining samples (Table 1; Fig. 8). Overall, the DNA viruses identified in the majority of samples collected during the Spring, Summer and Fall of 2021 were not found to be abundant (Table 1). The results of a Kruskal-Wallis Test indicated statistical differences (p–value = 1.45E-02) in the abundance of DNA viruses collected across seasons (Spring, Summer and Fall of 2021) with no differences (p–value = 2.00E-01) observed between locations in which samples were collected.

**FIG 8:**
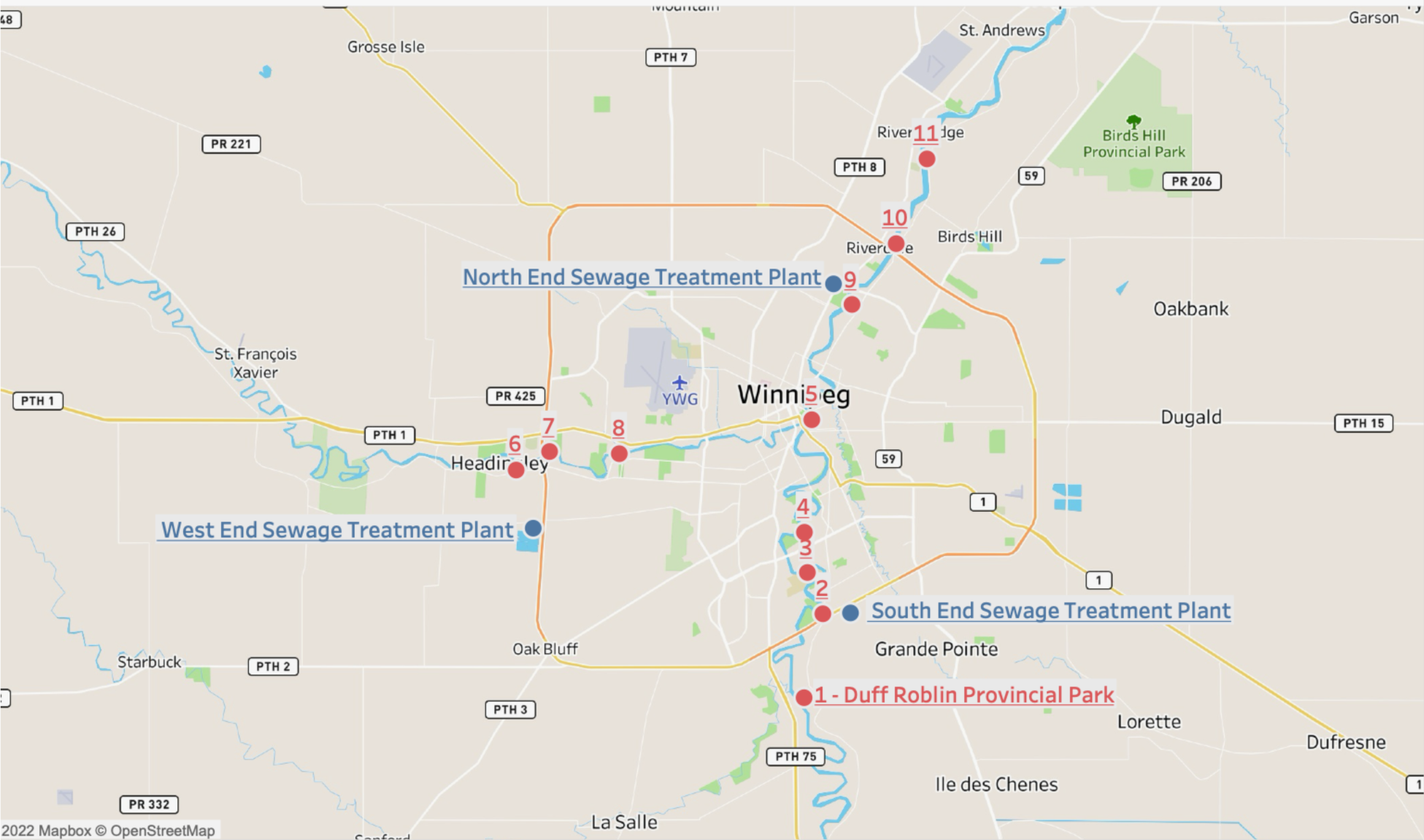
Map of 11 environmental sample locations indicated by the red dots. Winnipeg’s three major sewage treatment plants are indicated in blue as one of the sites visited even though no sampling events occurred. Source: Google maps and Tableau desktop (version 2022.3.1).

**Table 1:**
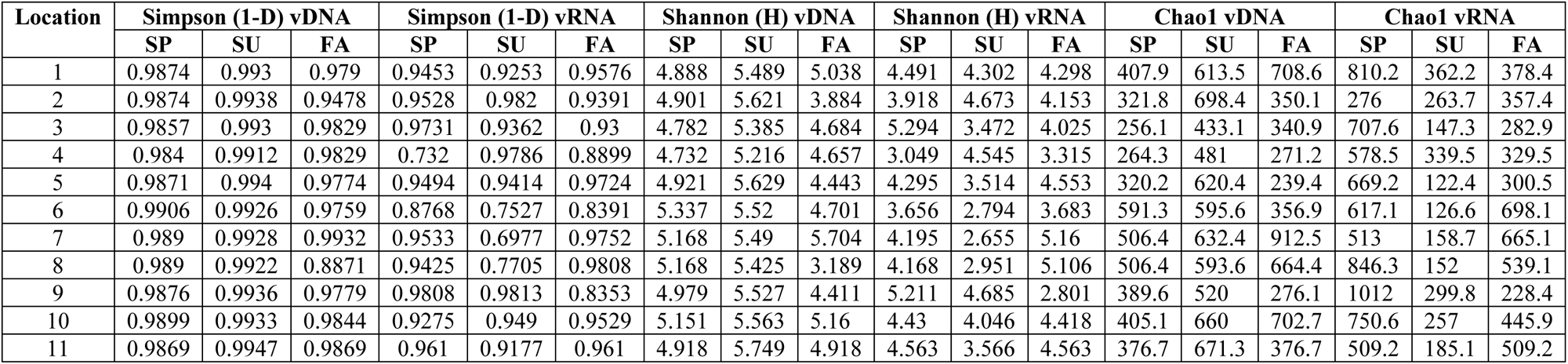
Assessment of Alpha Diversity Indices: Simpson (1-D), Shannon (H) and Chao1 among DNA and RNA viral communities present in samples 1-11 collected along the Red and Assiniboine Rivers of Winnipeg during the Spring, Summer and Fall of 2021. SP: Spring; SU: Summer; FA: Fall; vDNA: DNA viruses; vRNA: RNA viruses

**Table 2:**
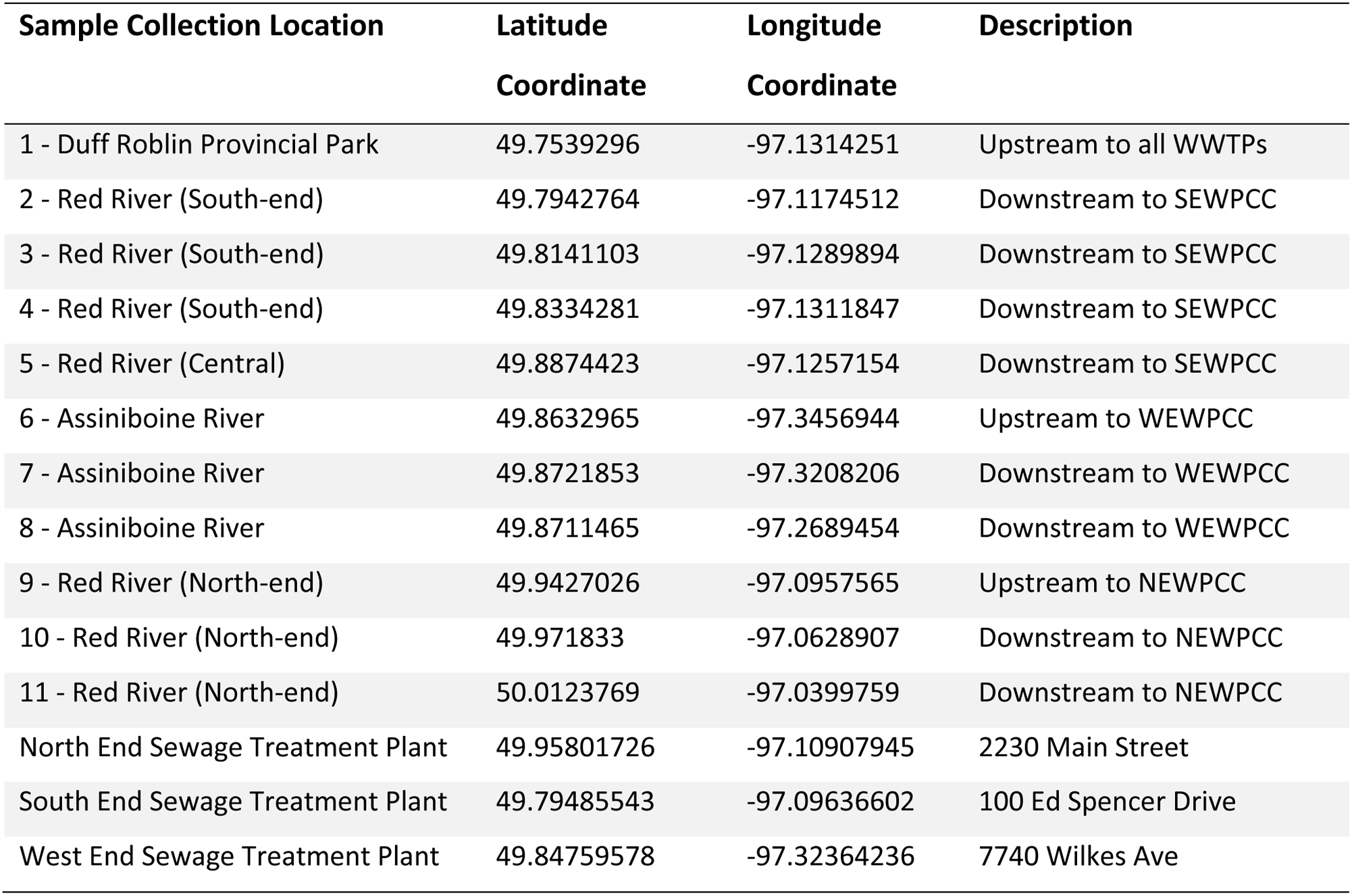
Coordinates for the 11 sample locations and Winnipeg’s three major sewage treatment plants.

| Sample Collection Location | Latitude<br>Coordinate | Longitude<br>Coordinate | Description |
| --- | --- | --- | --- |
| 1 - Duff Roblin Provincial Park | 49.7539296 | -97.1314251 | Upstream to all WWTPs |
| 2 - Red River (South-end) | 49.7942764 | -97.1174512 | Downstream to SEWPCC |
| 3 - Red River (South-end) | 49.8141103 | -97.1289894 | Downstream to SEWPCC |
| 4 - Red River (South-end) | 49.8334281 | -97.1311847 | Downstream to SEWPCC |
| 5 - Red River (Central) | 49.8874423 | -97.1257154 | Downstream to SEWPCC |
| 6 - Assiniboine River | 49.8632965 | -97.3456944 | Upstream to WEWPCC |
| 7 - Assiniboine River | 49.8721853 | -97.3208206 | Downstream to WEWPCC |
| 8 - Assiniboine River | 49.8711465 | -97.2689454 | Downstream to WEWPCC |
| 9 - Red River (North-end) | 49.9427026 | -97.0957565 | Upstream to NEWPCC |
| 10 - Red River (North-end) | 49.971833 | -97.0628907 | Downstream to NEWPCC |
| 11 - Red River (North-end) | 50.0123769 | -97.0399759 | Downstream to NEWPCC |
| North End Sewage Treatment Plant | 49.95801726 | -97.10907945 | 2230 Main Street |
| South End Sewage Treatment Plant | 49.79485543 | -97.09636602 | 100 Ed Spencer Drive |
| West End Sewage Treatment Plant | 49.84759578 | -97.32364236 | 7740 Wilkes Ave |

Whereas the DNA viral community evidenced consistently high species diversity amongst all surface water samples collected during the Spring and Summer of 2021 the diversity of RNA viral community fluctuated from sample to sample over the entire collection period (Table 1). Simpson’s (1-D) index suggests that RNA viruses in each sample were less diverse (Table 1) compared to that of DNA viruses (Table 1). Simpson’s (1-D) and Shannon diversity indices both suggest that low diversity of RNA viruses was present in surface water samples collected in the south end of the Red River (location 4; Table 2; Fig. 8) during Spring of 2021, in the Assiniboine River (locations 6, 7 and 8; Table 2; Fig. 8) during Summer of 2021, and in the north end of the Red River (location numbered 9; Table 2; Fig. 8) during Fall 2021 (Table 1). Although differences in species diversity of RNA viruses were not significant between sample locations (p–value ≥ 0.05), significant changes in RNA viral communities were observed across season (p–value < 1.0E-04 for Shannon index). However, Simpson’s (1-D) index suggests that species diversity across season was not significant (p–value = 7.46E-01). Compared to the other seasons, RNA viruses were observed to be most abundant in Spring and least abundant in the Summer of 2021 (Table 1). Similar to the relative abundances of DNA viruses, those of RNA viruses were also observed to be sparse. With overall lower values reported by Chao-1 richness index, it can be concluded that RNA viruses were less abundant possibly due to their low genome stability than that of DNA viruses among the samples collected. It can also be concluded that the relative abundances of DNA and RNA viruses identified were impacted by seasonality to a greater degree than the species diversities observed in each sample. Statistically, the abundance of RNA viruses observed differences (p–value < 1.0E-04) in samples collected seasonally, however no differences (p–value = 9.85E-01) in the abundance of RNA viruses were observed between sample locations.

## DISCUSSION

### Analyzing the Taxonomic Composition of DNA and RNA Viruses

DNA viruses maintain a high genomic stability due to their ability to use the host cell 3′-exonuclease proofreading activity and a variety of proteins as part of their reproductive process to perform checks and repair common kinds of DNA damage that will lower their mutation rates (Sanjuán et al., 2016; Duffy, 2018). In addition to the proofreading activity, many of the highly abundant DNA viruses identified in our study such as *Myoviridae, Podoviridae, Siphoviridae* and *Phycodnaviridae* are all non-enveloped in structure. Despite the absence of the lipid bilayer in “naked viruses”, they possess a very resistant protein capsid known to increase their survival to extreme environmental conditions and treatments such as heat, moisture, pH and chemical detergents (Firquet et al., 2015) and as such can resist changes in season. On the contrary, the distribution of RNA viruses was observed to be highly variable across season which may have led to even more inconsistencies in the taxonomy profiling across the viral databases used in the present study. Whereas the distribution of DNA viruses did not observe any major changes in seasons, the taxonomy of RNA viruses fluctuated across seasons with a large proportion remaining unclassified. Interestingly, NCBI RefSeq database (available at MG-RAST; Accession Number: PRJNA1011997) indicated that the control site consisted of 100 % unclassified viruses while the Kraken 2 Viral Genome database suggested that *Retroviridae* was 93 % predominant. As previously outlined, approximately 50 % of viral reads are classified as unknown when compared against viral databases (Goodacre et al., 2018; Fitzpatrick et al., 2021). Viral precipitation methods such as polyethylene glycol, iron chloride, ultrafiltration and skimmed-milk flocculation combined with extraction methods may be more biased towards DNA viruses rather than RNA viruses (Krishnamurthy et al., 2016; Hampton et al., 2020; Hjelmsø et al., 2017). This theory was confirmed in the present study which used skimmed-milk flocculation procedure as the primary method to concentrate viral particles from surface water samples. The findings from the Qubit 4 fluorometer which was used to quantify the total amount of extracted DNA and RNA from effluent samples (Table S1) showed significantly higher concentrations of viral DNA (ng/uL) across season, compared to viral RNA quantities (ng/uL) found to below the detection limit (< .025ng/uL) of the Qubit 4 fluorometer. Moreover, RNA viruses have smaller genome size ranging from 1.8 kbp up to 33 kbp and negatively correlated with mutation rate per site/replication (Duffy, 2018; Peck and Lauring, 2018; Caspi et al., 2023; Chaitanya, 2019; Campillo-Balderas et al., 2015). Enveloped viruses such as *Retroviridae* are typically very unstable because the lipid bilayer of the envelope is easily damaged by disinfectants and chemical detergents which often compromises the integrity of the virus to resist inactivation by harsh treatments and changes in weather (Firquet et al., 2015; Reitz and Gallo, 2015). These reasons may therefore allude to the observed variabilities of RNA viruses across season combined with random amplification method here used.

The taxonomy of DNA viruses identified from both NCBI RefSeq database (available at MG-RAST; Accession Number: PRJNA1011997) and Kraken 2 Viral Genome database were predominately DNA bacteriophages (*Myoviridae, Podoviridae* and *Siphoviridae*) belonging to the order *Caudovirales. Myoviridae, Podoviridae* and *Siphoviridae* collectively accounted for approximately 90 % of samples from each location along the Red and Assiniboine Rivers. With an approximate ubiquity ranging from 10^31^ – 10^32^, bacteriophages (phages) are regarded as the most abundant and highly dynamic entities in the biosphere (Hatfull and Hendrix, 2011; Kasman and Porter, 2022). Since the biology of DNA phages has been more studied, the majority of phages identified to date are tailed bacteriophages grouped under the order *Caudovirales*. In the same context, *Caudovirales* represented the largest order of bacteriophages and thus viruses here reported.

In comparison to the abundance of DNA phages, RNA phages (*Cystoviridae* and *Leviviridae*) were identified to a lesser extent accounting for approximately 3 % of samples collected at sites 5 and 7 in central Red River and Assiniboine River respectively during Fall 2021. Since RNA phages were only observed during the Fall of 2021 this may suggest that aquatic environmental conditions during the Fall of 2021 were the most favorable for RNA phages as opposed to DNA phages which appeared to survive well throughout season. Whereas DNA phages have been widely studied and successfully isolated from different aquatic environments very few studies have successfully isolated RNA phages from varying aquatic environments such that only two families *Cystoviridae* and *Leviviridae* of RNA phages have been discovered to date (Abd-Allah et al., 2021). In 2016 a study by Krishnamurthy *et al*. identified 122 partial genomes of RNA *Cystoviridae* phages from a transcriptome of pure culture of *Streptomyces avermitilis*. This publication was one of the first studies to identify RNA phages based on its affinity for a Gram-positive bacteria host which highlights the issue of many RNA phages that remain undiscovered in comparison to DNA phages (Krishnamurthy et al., 2016).

Majority of the RNA viruses (7%–100%) in the study locations were found to be *Retroviridae, Tombusviridae, Partiviridae*, *Picobirnaviridae*, Reoviridae*, Virgaviridae* (Fig. 3A; Fig. 3B). Of these top families, *Partiviridae, Tombusviridae, Picobirnaviridae*, *Reoviridae* and *Virgaviridae* are all non-enveloped or “naked viruses” which are highly resistant to wastewater treatment processes and environmental stressors due to their protein capsid that remains intact (Firquet et al., 2015). Although enveloped viruses such as *Retroviridae* are typically very unstable, *Retroviridae* has the survival advantage of genetic diversity as they have DNA intermediate in their replication cycle (Reitz and Gallo, 2015).

Interestingly, the Flavobacterium phage 11b was most abundant during the Spring of 2021 in the lesser polluted environment and at sites 5 and 10 in central and north end of the Red River respectively (Fig. S1) Bacteriophage 11b belongs to the order of Caudovirales and is known to infect the Flavobacterium found in cold, deep regions of aquatic environments with temperatures ranging from −20 °C to 20 °C (Borriss et al., 2007). In the present study, Flavobacterium phage 11b was identified during the Spring on May 16, 2021, when temperatures of the rivers were still cold from the winter period. Although the lesser polluted environment is situated upstream to the SEWPCC, the occurrence of Flavobacterium phage 11b in the lesser polluted environment may be due to the widespread problem of water pollution in which harmful chemicals, waste, plastic, and other pollutants promote the growth of pathogenic microorganisms in surface waters (Denchak, 2022).

Based on the results generated from PHASTER database, phages such as Pseudomonas phage UFV-P2, Rhizobium phage 16-3, Synechococcus phage S-CBS3, Thalassomonas phage BA3, Cyanophage MED4-117 and Ralstonia phage RSK1 DNA were only present in the Summer of 2021 (Fig. S1). This indicates that the composition of phages may be affected by changes in weather. Although no statistical analyses were performed in the present study to confirm seasonal variability of phages, many environmental studies have observed marked seasonal fluctuations of microbial communities in aquatic environments with low phage concentrations in the winter and high concentrations in the summer (Maurice et al., 2010; Chibani-Chennoufi et al., 2004; Vigneron et al., 2019). During the winter, there are reduced bacterial metabolisms, degeneration of organic matter and mobilization as well as a high proportion of damaged cells and as a result, lysogeny becomes the preferred strategy for phages to maintain their density (Vigneron et al., 2019). In northern Canada, lakes begin to change temperature at the beginning of Fall – November. Overall, the findings from PHASTER indicate that Spring 2021 harbored the greatest number of phages while only one phage-Rhodoferax phage P26218 was identified to be predominant in Fall 2021. It is possible that the weather conditions and factors during the Spring are more favorable for optimal survival for a diverse range of phages compared to Fall 2021. Furthermore, the findings from PHASTER are consistent with those phages identified by NCBI RefSeq database (available at MG-RAST) and Kraken 2 Viral Genome databases. Phages such as Vibrio phage, Thalassomonas phage BA3, Ralstonia phage, Synechococcus phage, Rhizobium phage 16-3, Pseudomonas phage UVF-P2, Myxococcus phage Mx8, Cyanophage and Flavobacterium phage 11b were identified from NCBI RefSeq database (available at MG-RAST) and Kraken 2 Viral Genome databases. Given the minority of RNA phages, Pseudomonas phage UVF-P2 was the only RNA phage identified from PHASTER while the remaining phages were DNA.

DNA viruses evidenced a consistently high abundance of approximately 40 % for the following functions: phage tail fiber proteins (5%–40%), phage replication (55%–38%) and phage packaging machinery (5%–33%) during the Spring, Summer and Fall of 2021 (Fig. 2). Phage tail fibers are attached to the bottom of the sheath vital in facilitating the attachment of the phage to the bacterium (Calero-Cáceres et al., 2019; Jebri et al., 2021). Following infection, phages initiate their ability to manipulate the metabolic and cellular machinery of the bacterium by halting the normal replication of bacterial components, and instead force the bacterium to replicate its own viral genome and structural components (Jebri et al, 2021; Call et al, 2019). Replication of the phage viral genome as part of the bacterium occurs via the lysogenic cycle if the host cell conditions are favorable, or via the lytic cycle if the host cell conditions are unfavorable. The phage then uses the bacterium’s machinery to synthesize additional proteins required to produce new phage particles (Davies et al., 2017; Marshall et al., 2021; Muniesa et al., 2013). Since the aquatic samples collected for the present study contained predominantly DNA phages (*Myoviridae, Podoviridae* and *Siphoviridae*), the phage-related functions determined from the MG-RAST SEED Subsystems (Fig. 2) may correspond to these group of DNA phages (Fig. 1; Fig. 3).

Given the high degree of variability among RNA viruses, it is no surprise that the functionalities of these viruses also differed across season, except for a few such as respiratory complexes, potassium homeostasis, and sialic acid metabolism that appeared to be consistent across season (Fig. 4). Although Fig. 4 depicts the top 20 functions of RNA viruses, approximately 300-400 functionalities were identified in the present study. Given the minority of RNA phages identified in aquatic samples collected for the present study, it is no surprise that the functionalities of RNA viruses were instead primarily related to metabolic pathways. Respiratory complexes are commonly found in respiratory RNA viruses such as *Respiratory Syncytial Virus, Influenza Virus, Parainfluenza, Metapneumovirus, Rhinovirus,* and *Coronavirus* and are therefore the major cause of acute upper and lower respiratory infections in young children, the elderly and the chronically ill (Curtsinger et al., 2023; Hodinka, 2016). Respiratory RNA viruses which maintain sialic acid metabolic functionality often have neuraminidase or a sialyl-O-acetyl-esterase and receptor binding proteins as part of viral envelopes or non-viral envelopes which allows them to bind to sialic acid receptors of humans as a point for viral entry to elicit an infection (Matrosovich et al., 2013). In summary, the most abundant functionalities of RNA viruses appear to most resemble the *Influenza virus* which may account for the abundance of cytochrome c oxidase (36%–52%) observed amongst RNA viruses in the Summer of 2021 (Fig.4). When overexpressed in H5N1 infected host cells, cytochrome c oxidase subunit 4 isoform 1 (COX41) has been found to be a positive regulator of Influenza A virus subtype H5N1 infection (He et al., 2022) by inducing apoptosis of the host’s native immune cells (Wang and Wang, 2022). The dissociation of cytochrome c oxidase into the cytoplasm is faciliated by PB1F2 membrane protein that is encoded by influenza virus genes to ensure a successful intracellular life cycle of the virus in host cells (Wang and Wang, 2022; Foo et al., 2022). Nevertheless, the taxonomic profiling from both viral databases (MG-RAST and Kraken 2) used for this study suggests that the Orthomyxoviridae family to which *Influenza virus* belongs to represents a minority of approximately 0.05 % of aquatic samples collected at sites 1 and 6 in the lesser polluted environments of the Red and Assiniboine river, during the Spring and Fall of 2021 respectively (*data not shown*).

### Evaluating Seasonal Variabilities of DNA and RNA viral communities

PMMoV and the DNA viruses (HAdV, crAssphage, *Myoviridae*, *Podoviridae*, *Herelleviridae*, *Herpesviridae* and *Phycodnaviridae*) were most influenced by Spring and Summer sampling events in the Red River as they were within the same quadrant as the Spring 2021 sampling event for sites 1–5 and 9–11 as well as the Summer 2021 sample collection sites 3 and 9–11along the Red River. On the contrary, the RNA viruses *Partitiviridae*, *Reoviridae*, *Betabaculovirus, Retroviridae*, *Picobirnaviridae* and *Peribunyaviridae* were observed within the same quadrant as the Summer and Fall 2021 sampling event for sites 6–8 along the Assiniboine River (Fig. 5; Table 2; Fig. 8). This indicates that the RNA viruses listed above were most influenced in the Assiniboine River during the Summer and Fall of 2021. Sampling conducted from areas of lesser polluted environments to those that were more influenced by anthropogenic activities including wastewater discharges (locations 1–5 and 9–11) was used to assess the influence of urbanization along the Red River. Thus, we propose that the abundance of DNA and RNA viruses were influenced by Factor 1 or “anthropogenic land use” where viral DNA communities in the Red River were influenced by urbanization and industrialization while the viral RNA communities in the Red and Assiniboine River were shaped by agricultural activities from upstream communities such as Portage la Prairie, Brandon, South Headingly and small rural municipalities.

Water quality parameters associated with the effect of urban influenced waterbodies include BOD_5_, cBOD_5_, TP, NH4-N and TN (Fig. 5; Ma et al., 2022). From our analysis, PMMoV was found to be influenced by the most water quality parameters associated with urbanization. To protect the health of residents relying on rivers for domestic and recreational usage, the City of Winnipeg WWTPs are required to obtain a mandatory wastewater license that stipulates routine daily tests to reduce levels of TN, BOD_5_, cBOD_5_, TP and NH4-N to improve the quality of effluents discharged. However, urbanization may also create environmental problems such as eutrophication and loss of biodiversity of aquatic species in receiving waterbodies (Flower et al., 2018). An increase in the population growth in Winnipeg is positively correlated to an increase in human enteric viruses PMMoV, HAdV and crAssphage excreted in human feces which enter the aquatic environments. For example, taxonomic classifications of assembled reads identified using the most recently updated Viral Genome Database revealed PMMoV belonging to the family *Tombusviridae* was highly abundant (53.1 %, 22.7 % and 75 %) in aquatic samples collected at sites 4, 5 and 9 respectively along the Red River (Fig. 3A; Table 2; Fig. 8).

We propose that the distinct clusters of sampling sites 6-8 in the Assiniboine River observed for the Spring, Summer and Fall sampling events (Fig. 5) contained viromes that were largely influenced by agricultural practices from communities situated upstream to the Assiniboine River. The Assiniboine River begins in eastern Saskatchewan and flows eastward through Brandon, Portage la Prairie and Headingley. Brandon, the second-largest city in Manitoba, has a large agricultural and manufacturing industry that processes flour, meat, fertilizers, chemicals, and petroleum products. The city Portage la Prairie is known for its diverse crop and livestock production capabilities due to its surrounding post-glacial flood plain that is highly fertile and rich with clay-loam soils abundant in nutrients. As of 2019, 3 % of a reported 3,000 jobs in the city Headingley area were agriculture, forestry, fishing and hunting, while 6 % were manufacturing. Although treatment processes at the City of Brandon Water Treatment Plant, Portage la Prairie’s Water Pollution Control Facility and Headingley Regional Water Treatment Plant may help to reduce the contaminants in industrial wastewater, we observed a close relationship in our exploratory analysis between the water quality parameters pH, TSS and DO and sampling locations 6-8 along the Assiniboine River during the Summer and Fall collection events (Fig. 5) which confirms the notion that these parameters are influenced by industrial and agricultural wastewater. For example, in Fig. 5 and Fig. 6 TSS was found to be positively correlated with *Peribunyaviridae* (r = 0.452; p = 8.20E-03) *Retroviridae* (r = 0.366; p = 3.63E-02) and *Picobirnaviridae* (r = 0.358; p = 4.09E-02). Agricultural pollution in receiving waterbodies is largely due to surface runoffs which increase turbidity and TSS levels. The development of farmlands loosens soil, increasing the opportunities for runoff and erosion. These runoffs often include nutrients such as nitrogen and phosphorous and pesticides commonly used on crops which subsequently promotes an influx in algae blooms and TSS levels in aquatic environments. Furthermore, high concentrations of TSS (mg/L) promotes the survival of waterborne microorganisms such as bacteria and viruses (Pinon and Vialette, 2018) that can withstand chemical disinfectants by protecting themselves in TSS aggregates.

In the present study, a weak but positive correlation was observed between the flow rate of river water and TSS (r = 0.303; p = 8.63E-02) (Fig. 5 and Fig. 6) indicating that an increase in flow rate subsequently increases the concentration of TSS. The effluent discharge was fastest (∼2.04 m/s^3^ and 1.82 m/s^3^) during the Spring 2021 sampling event at sites 1-5 and 9-11 along the Red River, which may disturb particulate matter such as clay, dirt, and soil from sediments settled along the riverbed and resuspend these within the waterbody as the river flows along its course. The Assiniboine River merges into the Red River at “The Forks” a central point along the Red River which is represented in our study as location 5. Therefore location 5 is subjected to an increase in flow rate as well as downstream locations 9-11 which receive a combined flow from the Assiniboine River and the southern end of the Red River. The flow rate of the Red River was found to have a significant positive influence on the enteric virus PMMoV assessed in the study (r = 0.375, p = 3.13E-02) (Fig. 5 and Fig. 6). This was further evidenced in models which explored the variability of factor 3 or “river water flow rate” (Fig. S2 and S3 in the supplemental material). Enteric viruses-HAdV, PMMoV, RoV and crAssphage are non-enveloped or “naked” viruses that are highly resistant to wastewater treatment processes and environmental stressors due to their protein capsid that remains intact (Firquet et al., 2015). Enteric viruses are therefore stable in high flows of water and can be transported to downstream locations (Derx et al., 2013). A Spearman’s correlation analysis further confirmed a significant positive correlation between HAdV and flow rate (r = 0.987, p-value = 3.03E-03) (Fig. 6) indicating that high river flows increase the transport of HAdV to downstream sites. Interestingly, water flow rate and DO appear to have the least impact on the viral community in our study as only the abundances of HAdV and *Mimiviridae* were significantly affected by flow rate and DO respectively (Fig. 5 and Fig. 6). Although water flow rate was not found to significantly influence the transport of most viromes identified, environmental studies have highlighted its importance (Gibson et al., 2011; Derx et al., 2013; Robins et al., 2019; Amoah et al., 2022). Although pH was observed to be within the same quadrant as sampling sites 6–8 along the Assiniboine River (Fig. 5) and is widely known to maintain a characteristic range in industrial wastewater, a Spearman correlation analysis (Fig. 6) revealed that pH was not significantly correlated to the abundances of viruses in the Red and Assiniboine Rivers. A possible reason for this may be due to the nature of RNA viruses identified in the Assiniboine River. *Reoviridae, Peribunyaviridae, Retroviridae* and *Betabaculovirus* were observed within the same quadrant as the Fall 2021 sample event for sites 6–8 (Fig. 5) are all enveloped viruses with highly resistant protein capsids to environmental conditions. During the Spring, Summer and Fall 2021 sampling events, the pH of the urban waterbodies was within normal range of 6.59-9.61 (Table S3) with more alkaline pH values observed at sites 6–8 in the Assiniboine River possibly due to agricultural runoffs into the river from upstream communities. Extreme fluctuations in pH values with values beyond the characteristic range of 6–9 in wastewater and effluents, may have inactivated the viruses present (Pinon and Vialette, 2018).

Factor 2 or “daylength and temperature” was the most influential factor on majority of the DNA and RNA viral communities identified in the study. Each of the enteric viruses – RoV, crAssphage and HAdV were found to have a significant positive correlation with daylength (r = 0.790, p < 1.00E-04) and temperature (r = 0.560, p = 6.00E-04) (Fig. 5). Furthermore, daylength and temperature were positively correlated with each other (r = 0.518, p = 2.00E-03) and as a result both factors exhibited a similar influence on the enteric viruses. High temperatures increase the viral inactivation rate due to the denaturing the secondary structures of proteins leading to impaired molecular viral function (Gamble et al., 2021; Pinon and Vialette, 2018). Studies which assessed the seasonality of RoV suggest that the biophysical state of proteins, lipids, and RNA molecules of the virus are affected by temperature (Bisht and Velthuis, 2022). In the present study, average temperatures of water collected across each season were range from 0 °C to 18 °C (Table S3). Although none of these temperatures may have significantly inactivated viruses (including enteric viruses), temperature plays a significant role in the composition of water microbiomes (Kim et al., 2017; Bertrand et al., 2012; Wyn-Jones and Sellwood, 2001). Daylength is another environmental factor known to increase viral inactivation. Sunlight emits infrared, visible, and ultraviolet (UV) rays. When UV light penetrates the cell walls of microorganisms, the molecular structure becomes permanently altered rendering them inactive and non-infectious. Although UV is effective at eliminating bacteria and parasites from wastewater at WWTPs, enteric viruses have displayed a degree of resistance to UV with HAdV being the most resistant enteric virus (Nwachuku et al., 2005; Rodríguez et al., 2022). In the present study, the daylength recorded for the Spring sample collection (May 16, 2021) was 15 hours while that for Summer and Fall sample collection (August 27 and November 21, 2021) were 13 hours and 8 hours respectively (Table S3). Since there was a gradual decrease in daylength across season, we propose that viral inactivation was highest during the Spring and lowest during the Fall of 2021. This was further confirmed in Fig. 5 which evidenced a close relationship was observed between daylength, temperature and the Spring and Summer sampling events.

Our beta diversity analysis indicated that the DNA viral communities in the Red River at locations 1–5 and 9–11 identified during the Spring of 2021 were found to be similar (Bray-Curtis dissimilarity distance of 2.71E-01) to those identified during the Fall of 2021 (Fig. 7A; Table 2; Fig. 8). This observational trend was also depicted in Fig. 5. Although Spring and Fall are seasons with contrasting characteristics, both consist of moderate temperatures ranging from −5°C to 20°C and high precipitation which may shape the microbial community in aquatic environments. Periods of heavy rains typically promote the spread of viral diseases transmitted by vectors such as Dengue virus, Yellow fever virus, West Nile Virus, Chikungunya Virus and Zika virus (Philip and Everlyn Polgreen, 2018), however, we have shown that in the present study precipitation has a significant influence and a negative correlation with HAdV (r = −0.735; p <1.00E-04) (Fig. 5; Fig. 6). As previously outlined, sampling occurred on May 16^th^ and November 21^st^, 2021, during the Spring and Fall. According to the historical data provided by the Government of Canada, Winnipeg experienced 54 mm of total precipitation (rainfall or snow) during the month of May 2021 and 37.2 mm during the month of November 2021 both of which were some of the highest amounts for the year. Furthermore, thunderstorm with light rain had occurred on May 15^th^ 2021, the preceding day of sample collection. Studies suggest that prolonged periods of rainfall may dilute pathogen concentrations in surface water (Kraay et al., 2020) as well as alter the viral concentrations from surface water runoff and resuspension of bottom sediments (Sassi et al., 2020; Shin et al., 2020). This rationale explains why our study observed a decrease in HAdV as precipitation increases.

The beta diversity measurement which indicates a diverse population of viruses in the Assiniboine River compared to that of the Red River during the Spring and Fall 2021 sampling events (Fig. 7A) is a similar pattern to the distinct clusters of sampling sites 6-8 from the Spring, Summer and Fall sampling events depicted from factor analysis (Fig. 5). As previously outlined, the effect of urbanization in Winnipeg may have shaped the viral community in the Red River while agricultural practices from communities such as Portage la Prairie, Brandon, and small rural communities influence the viral community in the Assiniboine River. Although observational trends of taxonomic classifications of assembled reads identified from NCBI RefSeq database (available at MG-RAST; Accession Number: PRJNA1011997) and Kraken 2 viral genome database revealed a relatively stable and consistent abundance of DNA viruses throughout changes in seasonal weather (Fig. 1), distance metrics analysis suggests that the taxonomic composition is most diverse during the Summer (Fig. 7A). A possible reason for the inconsistency in results may be due to the presence of additional viromes that were not previously identified from the metagenomic databases used within the study. A more refined update to the NCBI RefSeq database (available at MG-RAST) and Viral Genome database may improve the repository of newly sequenced organisms belonging to the viral community resulting in improved annotations based on similarity search and thus a difference in the relative abundances generated for this study. Given that the summer months are marked by warmer temperatures and increased sunlight, we would expect a reduction in the microbial community as these factors increase viral inactivation (Pinon and Vialette, 2018). However, the stability of DNA viruses may also increase resistance to environmental stressors.

RNA viral communities present in the Red and Assiniboine Rivers during the Fall appeared to be most similar and closely related (Bray-Curtis dissimilarity distance of 4.08E-02) to those identified in the Summer of 2021, while those present during the Spring were most diverse from the Summer and Fall sampling events (Fig. 7B).

Although RNA viruses were shown to be highly variable, observational trends depicted in Fig. 3 revealed a high similarity in the microbial community identified during the Summer and Fall sampling events with a high diversity for the Spring sample collection (Fig. 3). Although Summer and Fall appear to be characteristically contrasting seasons, the two maintain some similarities. As Summer transitions into Fall the warm temperatures observed during the Summer are also observed during the beginning of Fall before the temperature becomes cooler in later months. In our present study, temperatures during the summer sample collection ranged from 14.4 °C–19.4 °C while that during the Fall ranged between 0.9 °C–13.4 °C (Table S3). In addition, both Summer and Fall maintain similar lengths of days and night. As previously outlined, the average daylength during the Spring 2021 was 15 hours while that of Summer and Fall of 2021 were 13 hours and 8 hours respectively (Table S3). These reasons therefore allow for minimal seasonal change in the microbial diversity.

In comparison to RNA viral communities which were found to be more uniformed due to lower index values for Simpson’s (1-D) and Shannon (H) indices, DNA viruses measured high values for both indices which indicated high species diversity amongst all surface water samples collected (Table 1). While many RNA viruses are still yet to be discovered, DNA viruses have been more extensively studied and are found to contain considerable diversity especially among their genome structure. While RNA viruses maintain the advantage of high genetic diversity due to their increased susceptibility to mutation rates, DNA viruses maintain a higher species diversity than RNA viromes. This is because DNA viruses are classified with large genome sizes ranging from ≤ 2 kb up to 375 kb DNA which increases the diversity of organisms which DNA viruses can infect. (Payne, 2016). DNA viruses are known to infect organisms such as prokaryotes (bacteria and archaea) and eukaryotes (animals, plants, fungi, protozoans) (Campillo-Balderas et al., 2015; Harris and Hill, 2021). Whereas the genome sizes of DNA viruses vary by approximately four orders of magnitude, RNA viruses have more restricted genome sizes which vary up to one order of magnitude (Campillo-Balderas et al., 2015). To date, no reports or studies have indicated that RNA viruses are capable of infecting archaea (Bolduc et al., 2012; Campillo-Balderas et al., 2015) and one possible reason for this may be due to their limited genome sizes coupled with the fact that many RNA viruses remain undiscovered. In the present study, significant changes in RNA viral communities were observed across season (p–value < 1.0E-04 for Shannon index), however, Simpson’s (1-D) index suggests that species diversity across season was not significant (p–value = 7.46E-01) (Table 1). The variability in RNA viral communities is strongly subjective to the type of diversity index used. Simpson (1-D) index is biased towards the most dominant or relative abundant species, while Shannon Index considers all species which are represented in a sample and is strongly influenced number of species, i.e. richness and the average or evenness of individual distribution in the species. Overall, our findings display a higher species diversity for DNA viruses than RNA viruses thereby confirm current literature.

In the present study, we observed highest species diversity and richness of DNA viral communities during the Spring and Summer 2021 compared to Fall 2021 (Table 1). Viruses typically display marked seasonal peaks although they may be present throughout the year. It is possible that the conditions of Spring and Summer may be favorable to host diverse DNA viral communities in the Red and Assiniboine River. Overall, DNA viruses were found to be more abundant than that of RNA viruses among aquatic samples collected in our study possibly because DNA viruses maintain a high genomic stability with low mutation rates (Sanjuán et al., 2016; Duffy, 2018). Additionally, many of the highly abundant DNA viruses identified in our study such as *Myoviridae, Podoviridae, Siphoviridae* and *Phycodnaviridae* are all non-enveloped in structure which increases their survival to extreme environmental conditions and treatments such as heat, moisture, pH and chemical detergents (Firquet et al., 2015). In conclusion, our findings on the index values for Simpson’s (1-D) and Shannon (H) indices indicate that the relative abundances of DNA and RNA viruses identified were impacted by seasonality to a greater degree than the species diversities observed in each sample. However, viral communities can be affected by several factors. As previously mentioned, our findings from EFA suggest that viral RNA and DNA communities in the Red and Assiniboine Rivers were further influenced by urbanization, industrialization and agricultural activities in Manitoba.

Future directions of this research project require continuous sequencing and validation methods over longer sampling event periods combined with longer sequencing technologies to accurately establish the characteristics and dynamics of the viral community structures in aquatic environments influenced by anthropogenic activities. This may ultimately lead to the development of novel viral markers of fecal contamination. Although it is undeniable the role that WWTPs have played in maintaining water quality, they are not designed to remove all genetic material or microbes. In this context, screening of viruses is not a common practice to assess efficiency of wastewater treatment. We hope to collaborate with operators at wastewater treatment facilities, local governments, civil engineers, and/or health policymakers to include viral screening procedures as part of routine water quality testing. Although all available viral wastewater treatments pose major drawbacks, we propose the use of ozonation and/or Membrane BioReactors (MBR) followed by biological post-treatments as promising methods to reduce viral loads to go into the environment. Ozonation of sewage has been found to further reduce the environmental spread of enteric viruses-Adenovirus, Norovirus, Astrovirus as well as bacteria and bacteriophages by one to two log_10_ during quantitative PCR analysis (Wang et al., 2018), while the use of MBR has led to has led to an overall decrease in viruses from 52.03% to 15.04 % including a reduced proportion of MS2 bacteriophages (Zhang et al., 2022).

By determining the differences between environmental samples’ fingerprints and effluents from WWTPs, we will be able to identify the most informative viral genes associated these discharges necessary to implement appropriate methods to screen and remove pathogenic viruses during municipal wastewater treatment. Furthermore, fecal indicator bacteria combined with the detection of enteric viruses may complement and improve assessment of water quality in effluents discarded into rivers.

## Materials and Methods

### Surface water sampling

To evaluate seasonal variabilities, aquatic sampling was conducted at 11 locations along the Red and Assiniboine Rivers in the Spring on May 16^th^, Summer on August 27^th^ and Fall on November 21^st^ 2021. Of the 11 locations, nine of them were located on the proximity or downstream of the discharge points from Winnipeg’s three major sewage plants. Two sample collection sites were used to represent a case-control study from undisturbed or lesser polluted environments. Location 1 was upstream of the NEWPCC outfall in the Red River situated outside the Winnipeg Metropolitan area (Duff Roblin Provincial Park) while location 6 was situated upstream of the WEWPCC outfall in the Assiniboine River (Fig. 8; Table 2). As the numbers increase from the control sites, they are located further away from each of the wastewater treatment plants. In addition, MilliQ water was used as a negative background control. A total of thirty-six 10-L surface water samples were analyzed over the three sampling event periods. All samples collected were stored in a 4 °C cold room until processing occurred within 24 hours following sample collection.

### Concentrating viruses from environmental samples

To minimize noise during metagenomic sequencing from larger fractions such as micro-eukaryotes and bacteria to the smaller ones such as viruses, raw water samples were first subjected to capsule and vacuum filtration using PALL Corporation GWV high-capacity groundwater sampling capsules of 0.45 µm size and PALL membrane disc filters of 0.2 µm and 0.1µm sizes. The filtrate then underwent SMF where the skimmed-milk particulates target the non-settling particles such as viruses and forces it to clump together into larger and heavier solids known as “flocs” by means of sedimentation. A modified SMF approach described by Calgua et al. (2008) and Fernandez-Cassi et al. (2017) was used to concentrate virus particles (Yanaç et al., 2023). A 1 % wight-by-volume pre-flocculated skimmed milk solution was prepared using 1.32-L of RICCA synthetic seawater (ASTM D 1141 substitute ocean water without heavy metals) and 13.2 g of Difco skimmed milk powder and then acidified to a pH of 3.5 using 2 M HCL. 100 mL of the pre-flocculated acidified skimmed milk solution was transferred to 10-L acidified (pH 3.5) environmental water samples [final skimmed milk concentration 0.01 % (w/v)]. Using a magnetic stirrer and a magnetic stir bar, samples were stirred for 8 hours at room temperature and allowed to settle down by gravity for an additional 8 hours. Without disturbing the flocs, the supernatant was carefully removed using a vacuum pump. The remaining flocs was aliquoted and balanced according to weight in 50 mL centrifuge tubes for centrifugation at 8000 g for 30 mins at 4 °C. The resulting pellet which theoretically contained viruses of interest, was dissolved in 0.2 M phosphate buffer. To eliminate free DNA and RNA present that may be co-precipitated into the kit used for nucleic acid extraction, the dissolved pellet was treated with DNAse I, and RNAse A (Thermo Fisher Scientific, Waltham, MA, USA). As outlined by the manufacturer (Thermo Fisher Scientific), 3 µL of 10 mg/µL RNAse A was added to the dissolved pellet and incubated at 37 °C for 30 minutes. For DNAse treatment, 30 µL of 10X reaction buffer with MgCl_2_ and 3 µL of 1 unit/µL DNAse I were added to pellet and incubated at 37 °C for 10 minutes. To inactivate the DNAse enzyme, 3 µL of 50mM EDTA was added and incubated at 65 °C for 10 minutes. Inactivation of RNAse A enzyme occurred though nucleic acid extractions.

### Extracting and Quantifying Total Nucleic Acids

A Qiagen All-Prep DNA/RNA power microbiome kit was used to extract total nucleic acids from the dissolved pellet. As outlined by Qiagen quick-start protocol (2017), 600 µL of warmed solution of PM1, 450 µL effluent sample, 100 µL ultrapure phenol:chloroform:isoamyl alcohol (Invitrogen, Thermo Fisher Scientific) and 6 µL 2-mercaptoethanol (Fisher Chemical, Fisher Scientific) were added to a Qiagen powerbead tube. Using a vortex adapter, the powerbead tubes were vortexed horizontally at maximum speed for 10 minutes and then centrifuged at 13 000 g for one minute at 22 °C. The supernatant was transferred to a clean 2 mL Collection Tube where 150 µL of IRS Solution was added and incubated at 4 °C for five minutes. The 2 mL collection tubes were centrifuged at 13 000 g for one minute. The flow through was discarded and ∼700 µL of the supernatant was transferred to clean 2.2 mL collection tubes. To each collection tube, 600 µL solutions PM3 and PM4 were added and vortexed briefly. For each effluent sample, 625 µL of the supernatant was loaded into a MB spin column and centrifuged at 13 000 g for one minute. The flow through was discarded and the process was repeated until the all the supernatant was loaded into the MB spin columns for each sample. To wash the membrane of the MB spin column which in theory contains the extracted nucleic acids, 600 µL of solution PM4 was added and centrifuged at 13,000 g for one minute. The flow through was discarded and the process was repeated with 600 µL of solution PM5 that was added to the membrane. To dry the membrane of the spin column, the empty column was centrifuged at 13,000 g for two additional minutes. To elute the total nucleic acids from the membrane of the spin column, 100 µL of warmed RNAse-free water was added to the centre of the white column membrane and incubated for two minutes. The MB spin column was then centrifuged at 13,000 g for one minute. The final volume of extracted nucleic acids (100 µL) was transferred to a clean collection tube and used for downstream applications. A Qubit 4 fluorometer (Invitrogen, Carlsbad, CA, USA) was used to quantify the total amount of extracted DNA and RNA from effluent samples.

### Enriching Viral DNA and RNA Fractions

To enrich viral RNA and viral DNA fractions and avoid the degradation of RNA which is less stable than DNA and prone to being degraded by exogenous ribonucleases, the final volume of the total extracted nucleic acids (100 µL) was separated into half. For each sample, 50 µL were treated with Turbo DNAse (Thermo Fisher Scientific, Waltham, MA, USA) and the other half (50 µL) was treated with Ambion RNAse III (Thermo Fisher Scientific, Waltham, MA, USA). As outlined by the manufacturer’s protocol, 1 µL of 1 unit/µL RNAse III and 5µL of 10X RNAse III reaction buffer was added to the viral DNA fraction and incubated at 37 °C for one hour. To inactivate the RNAse III, the DNA fraction was incubated at 95 °C for 5 minutes. To reanneal the DNA, the sample tubes were placed on ice for 8 minutes. As indicated by the manufacturer’s protocol, 2 µL of 2 units/µL Turbo DNAse and 5 µL of 10X Turbo DNAse buffer was added to the viral RNA fraction and incubated at 37 °C for 30 minutes. To inactivate the Turbo DNAse, 5.5 µL of homogenized DNAse inactivation reagent in a 1:10 proportion was added to the viral RNA fraction and allowed to centrifuge at 10,000 g for 1.5 minutes at 22 °C.

The treated total RNA fraction contained viral RNA, messenger RNA (mRNA), small RNAs and ribosomal RNA (rRNA). Therefore, a QIAseq FastSelect kit (Qiagen Sciences, Maryland, MD, USA) was used for rapid removal of bacterial and mammalian rRNA. As outlined by the QIAseq FastSelect 5S/16S/23S rRNA handbook from Qiagen, 1.5 µL of 12 µL FastSelect FH Buffer, 1µL from 8µL Fast select 5S/16S/23S enzyme previously incubated at 37 °C for 5 minutes, 1 µL of 12 µL Fast Select rRNA HMR enzyme, and 1.5 µL of nuclease-free water were all added to 11 µL of viral RNA fraction and incubated in a MiniAmp thermal cycler (Applied Biosystems, Waltham, MA, USA) for 5.5 minutes at 89 °C, 2 minutes at 75 °C, 2 minutes at 70 °C, 2 minutes at 65 °C, 2 minutes at 60 °C, 2 minutes at 55 °C, 2 minutes at 37 °C, 2 minutes at 25 °C and then cooled down at 4 °C. Homogenized QIAseq beads (19.5 µL) were added to the total volume vortexed and incubated for 5 minutes at room temperature. To separate the bacterial and mammalian rRNA from the QIAseq beads containing the RNA of interest, the tubes were placed on a magnetic rack for 2 minutes until the solution became clear and the beads were on the magnetic rack. The supernatant was carefully discarded without disturbing the QIAseq beads. 1.5 µL of nuclease-free water and 19.5 µL of 10.2 mL QIAseq bead binding buffer was added to the QIAseq Beads, vortexed and incubated for 5 minutes at room temperature. The tubes were placed on a magnetic rack for 2 minutes until the solution cleared and the beads were on the magnetic rack. The supernatant was carefully discarded without disturbing the QIAseq beads containing the RNA of interest. To wash the QIAseq beads, 200 µL of 80 % ethanol was added to the tubes, and the supernatant was discarded. The QIAseq beads were air dried for 2 minutes until all the residual supernatant had evaporated. To elute the viral RNA fraction of interest, 31 µL of nuclease-free water was added to the QIAseq beads. The supernatant was vortexed and incubated at room temperature for 2 minutes to appropriately hydrate the QIAseq beads. The tubes were placed on a magnetic rack for two minutes until the solution cleared and the beads were on the magnetic rack. The supernatant containing the RNA of interest had a final volume of ∼29 µL. To obtain quantifiable amounts of viral RNA necessary for sequencing, whole transcriptome amplification was conducted using a REPLI-g cell WGA & WTA kit that enables amplification of viral DNA and RNA in parallel reactions. For DNA viral fractions, this step was not required as quantifiable amounts were observed after the addition of Ambion RNAse III enzyme to degrade ribosomal rRNA.

As outlined by the REPLI-g cell WGA & WTA handbook from Qiagen, 8 µL of lysis buffer was added to 13 µL of the final RNA fraction of interest and was incubated at 24 °C for 5 minutes followed by 95 °C for 3 mins and cooled down at 4 °C. To the lysed cell sample, 2 µL of gDNA wipeout buffer was added and incubated at 42 °C for 10 minutes to ensure gDNA removal. Eight µL of Quantiscript RT mix was freshly prepared using 4 µL RT/polymerase buffer, 2 µL H_2_O, 1 µL of 0.4 µg/µL oligo dT primer and 1 µL Quantiscript RT Enzyme Primer for a single reaction. Eight µL Quantiscript RT Mix was then added to 23 µL of the lysed and unsheared cell sample (as described above), briefly vortexed and incubated at 42 °C for 60 minutes to allow for reverse transcription. To stop the reverse transcription reaction, the sample was incubated at 95 °C for 3 minutes and cooled down at 4 °C. Ten µL of freshly prepared ligation mix was prepared from 8 µL 10X ligase buffer and 2 µL ligase mix for a single reaction. 10 µL ligation mix was added to 31 µL of the Quantiscript RT reaction, vortexed briefly and incubated at 24 °C for 30 minutes to allow for ligation. To stop the ligation reaction, the sample was incubated at 95 °C for 5 minutes and cooled down at 4 °C. Thirty µL of REPLI-g SensiPhi amplification mix was freshly prepared from 29 µL REPLI-g Single Cell Reaction Buffer and 1 µL REPLI-g SensiPhi DNA Polymerase for a single reaction. Thirty µL of REPLI-g SensiPhi amplification mix was added to 41 µL of the ligation reaction, vortexed briefly and incubated at 30 °C for 2 hours to randomly amplify the cDNA. To inactivate the DNA polymerase, the reaction was incubated at 65 °C for 5 minutes and cooled down at 4 °C. The final volume of randomly amplified cDNA (71 µL) was used to establish a baseline of virome distribution through metagenomics and preliminary screening of enteric viruses.

### Characterizing Virome Distribution

Metagenomics was applied to explore and characterize the DNA and RNA virome distribution in urban influenced environments. Enteric viral markers quantitatively assessed in each effluent sample were: *Human Adenovirus 40/41* (HAdV), *Cross Assembly Phage* (crAssphage), *Pepper mild mottle virus* (PMMoV) and *Rotavirus* (RoV), as previous literature and results in the Uyaguari laboratory has evidenced these enteric viruses to be relatively more stable enteric viral markers of fecal contamination (Garcia et al., 2022). Using the results generated from these approaches, seasonal patterns of viral communities were then determined using exploratory factor analysis, distance metrics on the beta diversity index, and generalized linear model (GLM) on the alpha diversity indices.

### Metagenomic Analyses

A total of 36 10-L environmental samples collected over the three sampling event periods once nucleic acid extracted were prepared for sequencing. Fifteen µL of viral DNA and randomly amplified RNA fractions were sent to Integrated Microbiome Resource (IMR) for Illumina sequencing (Dalhousie University, Halifax, NS, Canada; Comeau et al., 2017). A mock community of pooled DNA and RNA viruses was also included to account for metagenomic sequencing controls. DNA viruses included *Adenoviruses* (such as *Adenovirus viral soup extract* and *Adenovirus 1*) and *Myoviruses* such as *Myophage g20(+) M2 and Myophage g20(+) M3*. RNA viruses included *Enteroviruses* (such as *Enterovirus* and *Enterovirus Coxsackie B2*) and *Heterosigma akashiwo* RNA virus (HaRNAV). These viruses adhered identically to the protocol used to concentrate human enteric viruses from environmental water samples and were then pooled in equimolar concentrations for metagenomic sequencing at IMR (Dalhousie University, Halifax, NS, Canada; Comeau et al., 2017). As outlined by IMR, samples sent for sequencing were tagmented, amplified through PCR techniques, barcoded for identification, purified using columns or beads and normalized using Illumina beads. To generate paired-end Illumina sequenced reads, a pooled library was prepared using an Illumina Nextera Flex kit and ∼1 ng of DNA samples (Comeau et al., 2017; IMR, 2014; Peterson et al., 2021). Massive parallel sequencing was conducted using NextSeq 550 System (Illumina Technology) and NextSeq 550 System High-Output Kit to produce ∼4 M Pair Ended reads and 2.4 Gb/sample sequence (Comeau et al., 2017; IMR, 2014; Peterson et al., 2021).

Geneious Prime bioinformatic platform was used to pre-process the NGS reads necessary to facilitate identification of virome distribution by Metagenomics Rapid Annotation using Subsystem Technology (MG-RAST) server (Meyer et al., 2008). Preprocessing of high-throughput reads adhered to the Geneious Prime alignment and assembly workflow which provided steps to enhance accuracy and reduce computation time required to assemble the raw reads (Gibbs, 2020; Geneious Prime, 2022; Kearse et al., 2012). The paired read data provided by IMR as separate forward and reverse compressed gzip sequences in fastq format was imported into Geneious Prime. Using the set paired reads operation from the BBMerge function accessible through the pre-processing sequence menu, the forward and reverse sequences were paired for each sample. The raw reads were not trimmed in Geneious Prime as the adapter trimming quality control step was completed by IMR using Nextera Flex DNA prep. Duplicate, contained and overlapping sequences were then removed from the dataset using the Dedupe function containing dedupe python library and machine learning algorithms, available from the remove duplicate reads option in the sequence menu (Geneious Prime, 2022; Gibbs, 2020). Using BBMerge function as part of the BBtools suite found in the merge paired reads option in the sequence menu, two overlapping paired end reads were joined together by overlapping detection to form a single read (Geneious Prime, 2022; Gibbs, 2020). To down-sample reads in high-depth areas of a genome and eliminate false-positive variant calls, the reads were normalized and corrected for error using the K-mer based bioinformatic function BBNorm, a member of the BBTools package (Brian Bushnell, 2014; Geneious Prime, 2022; Gibbs, 2020). The pre-processed paired end reads were assembled and contigs were generated using the De novo assembly method within Geneious, Prime.

To facilitate in depth viral metagenomics of sequenced results, Kraken 2 taxonomic sequence classification system and MG-RAST metagenomic analysis server were used to identify assembled reads as DNA and RNA viruses. The contigs (assembled pre-processed paired end reads) from Geneious Prime bioinformatic software platform was uploaded into MG-RAST version 3.6 (Meyer et al., 2008). Taxonomic classifications of assembled reads as DNA and RNA viruses were identified using the RefSeq: NCBI reference sequences database at the species level. Hierarchical functionalities of assembled reads as DNA and RNA viruses were identified using the subsystem database at level three.

To establish a comparison of taxonomic classifications across viral databases, the raw sequenced reads were also uploaded into Kraken 2 taxonomic sequence classification system which relies on an assembly-free method (Wood et al., 2014). Kraken 2 taxonomic sequence classification system was accessed through Galaxy Europe, a graphical user interface with powerful data analytical tools to facilitate a wide range of bioinformatic analyses. Contigs were first generated from the raw sequenced reads as separate forward and reverse in fastaq format using the *Make.contigs* tool (Schloss et al., 2009) accessible through Galaxy Europe (The Galaxy Community, 2022). The contigs were then uploaded into Kraken 2 taxonomic sequence classification system as single reads. Taxonomic classifications of assembled reads as DNA and RNA viruses were identified using the most recently updated viral genome database (2019). To avoid false positive results, the confidence score threshold was set to 0.5 as a recommended cut off point for low quality hits (Kraken operating manual, 2022). Taxonomic classifications were generated at the family level. The Kraken 2 and MG-RAST reports generated for each sample were compiled together and visualized in Tableau desktop (Tableau version 2022.3.1) to depict the relative abundance (%) across each sample location and thus seasonal variabilities.

### Quantitative Analyses

Enzymatic treatments of nucleic acid extracts enriched for viral DNA were quantitatively assessed through real time quantitative polymerase chain reaction (qPCR) to identify the DNA viruses - HAdV and crAssphage. As described by the CDC, 2020 and Corman et al, 2020, a 10 µL targeted 2.0d qPCR reaction for each DNA enteric virus contained 5 µL of Taqman environmental master mix, 500 nM of each specific primers-AdV-F, AdV-R, 056F1 and 056R1, and 250 nM of probes Adv-P and 056P1 (Table S2; Stachler et al., 2018; Molecular Microbiology & Genomics Team, British Columbia Centre for Disease Control, 2017). To generate a standard curve necessary for quantifying DNA, a targeted synthetic oligonucleotide such as a gBlock Gene Fragment was used for each assay. Seven positive controls/standards were made from serial dilutions of Adeno/Crass gBlock at 7.06×10^7^ copies/µL. A non-template control consisting of nuclease free-water (Promega Corporation, Fitchburg, WI, United States) was used for each assay. HAdV and crAssphage were subjected to the following targeted qPCR conditions: 50 °C for 2 min and 95 °C for 10 min followed by 40 cycles of 95 °C for 15 s and 60 °C for 1 min. Nucleic acid extracts enriched for viral RNA with no random amplification step were assessed through reverse transcription quantitative real-time PCR (RT–qPCR) to quantify PMMoV and RoV. A 10 µL RT–qPCR reaction to identify the RNA enteric viruses of interest contained 2.5 µL of fast virus 1-step master mix (4X) (Life Technologies, Carlsbad, CA, USA), 500 nM of each specific primers: PMMV-FP1-rev, PMMV-RP1, NSP3-F, and NSP3-R, and 250 nM of each probe: PMMV-P and NSP3-P (Table S2; Zheng et al., 2008; Rosario et al., 2009). PMMoV and RoV were subjected to the following RT–qPCR conditions: 50 °C for 5 min and 95 °C for 20 s followed by 40 cycles of 95 °C for 3 s and 60 °C for 30 s.

A QuantStudio 5 real-time PCR system (Applied Biosystems, Waltham, MA, USA) was used to conduct quantification of enteric viruses in thirty-six, 10-L environmental samples collected over the three sampling event periods. All qPCR and RT–qPCR reactions used 2 µL of template. Each sample screened for an enteric virus of interest, all positive controls and non-template controls (NTC) were run in triplicates. For a sample to be considered as positive, 2 out of 3 of its replicates had to be amplifiable. According to the Public Health Ontario the cut-off (highest Ct value) for positivity of a sample is at 38 cycles (Public Health Ontario, 2020). Thus, those above 38 cycles (from a total of 40 cycles) were considered as a negative result. Raw output data was analyzed following the methodology described by Ritalahti et al. (2006). Gene copy numbers (GCN) per volume were determined from the following equation:

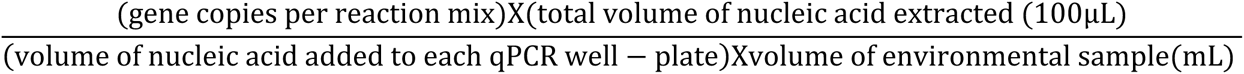

### Statistical analyses to determine Seasonal Patterns of viral communities

To explore the strength of the hypothesized relationships between viral families and latent environmental conditions at each sample site across season, an exploratory factor analyses (EFA) was conducted in Statistical Analysis System (SAS, version 9.4 for Windows) and visualized using Tableau desktop (Tableau version 2022.3.1). The following logarithmic physicochemical and biological parameters were included in the EFA model: (i) <u>publicly available metadata</u> from Winnipeg’s WWTPs such as TSS (mg/L), BOD_5_ (mg/L), cBOD_5_ (mg/L), NH4-N (mg/L), Ortho-Phosphorous (mg/L-P), TP (mg/L), E-coli (MPN/mL) and TN (mg/L), (ii) <u>environmental metadata</u> obtained using a YSI probe such as temperature (°C), pressure (mmHg), Salinity (psu) and dissolved oxygen levels (mg/L), and (iii) <u>information from meteorological stations</u> in Winnipeg, MB located nearby each sampling site, such as cumulative precipitation (mm), daylight length (mins) and discharge of water flow rate (primary sensor derived (m^3^/s)). The characteristics of effluent wastewater samples for the dates the sample were collected are summarized in Appendix A.1. Orthomax rotation that best fits all variables assessed in this study was used on the EFA model. A scatter plot was generated in order to visualize the factor loadings for the environmental parameters in relation to each sample site and season. A correlogram was also generated to assess the strength of the relationship between the water quality parameters and the relative abundance of the top ten viral DNA and RNA communities identified metagenomically alongside the gene copy numbers of enteric viruses assessed quantitatively.

Using the Paleontological Statistics Software Package for Education data analyzer software (PAST 4.3, Hammer et al., 2001), the richness (Chao-1), alpha diversity indices such as Simpson (1-D) and Shannon (H), as well as the Bray-Curtis dissimilarity (Beta Diversity) index were calculated from the outputs of the taxonomic classifications generated by MG-RAST server. Alpha and Beta diversities were then visualized in Tableau desktop (Tableau version 2022.3.1) as tables and scatter plots respectively to depict seasonal variabilities. To determine statistical differences of each alpha diversity indices as well as Chao-1 richness between sample locations and between seasons of each sample collection period, a Kruskal-Wallis test and multiple comparisons were conducted using the NPAR1WAY procedure in the Statistical Analysis System (SAS, version 9.4 for Windows). To determine the similarities and dissimilarities between samples collected during the Spring, Summer and Fall 2021, the Bray-Curtis distance matrix from the *Abdiv* package in R and R studio (RStudio Team, 2021; R Core Team, 2021) was calculated.

## Acknowledgements

The authors would like to acknowledge the following research organizations for grants used to procure reagents necessary to conduct in-laboratory research: Visual Automated Disease Analytics (VADA) graduate training program, Mitacs for granting an Accelerate Fellowship (IT26939), the Natural Sciences and Engineering Research Council of Canada for the grants: NSERC-Alliance Grant No. ALLRP 553987 and NSERC-Discovery Grant No. RGPIN-2022-04508), as well as the research start-up funds (grant No. 322388). We would like to acknowledge that the University of Manitoba campuses are located on the original lands of the Anishinaabeg, Cree, Ojibwe-Cree, Dakota, and Dene peoples, and on the homeland of the Métis Nation.

## Supplemental Material

**Table S1:**
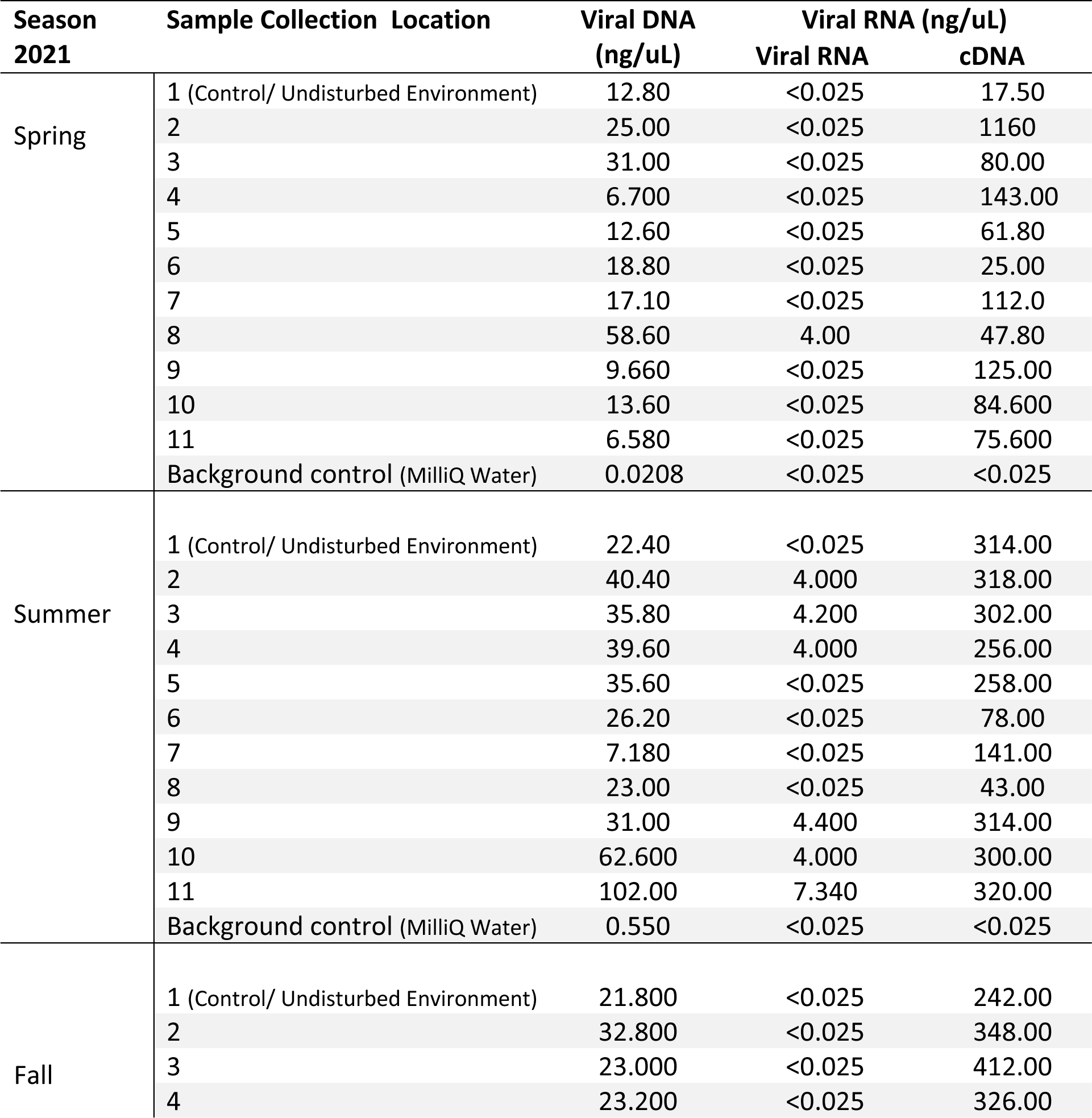
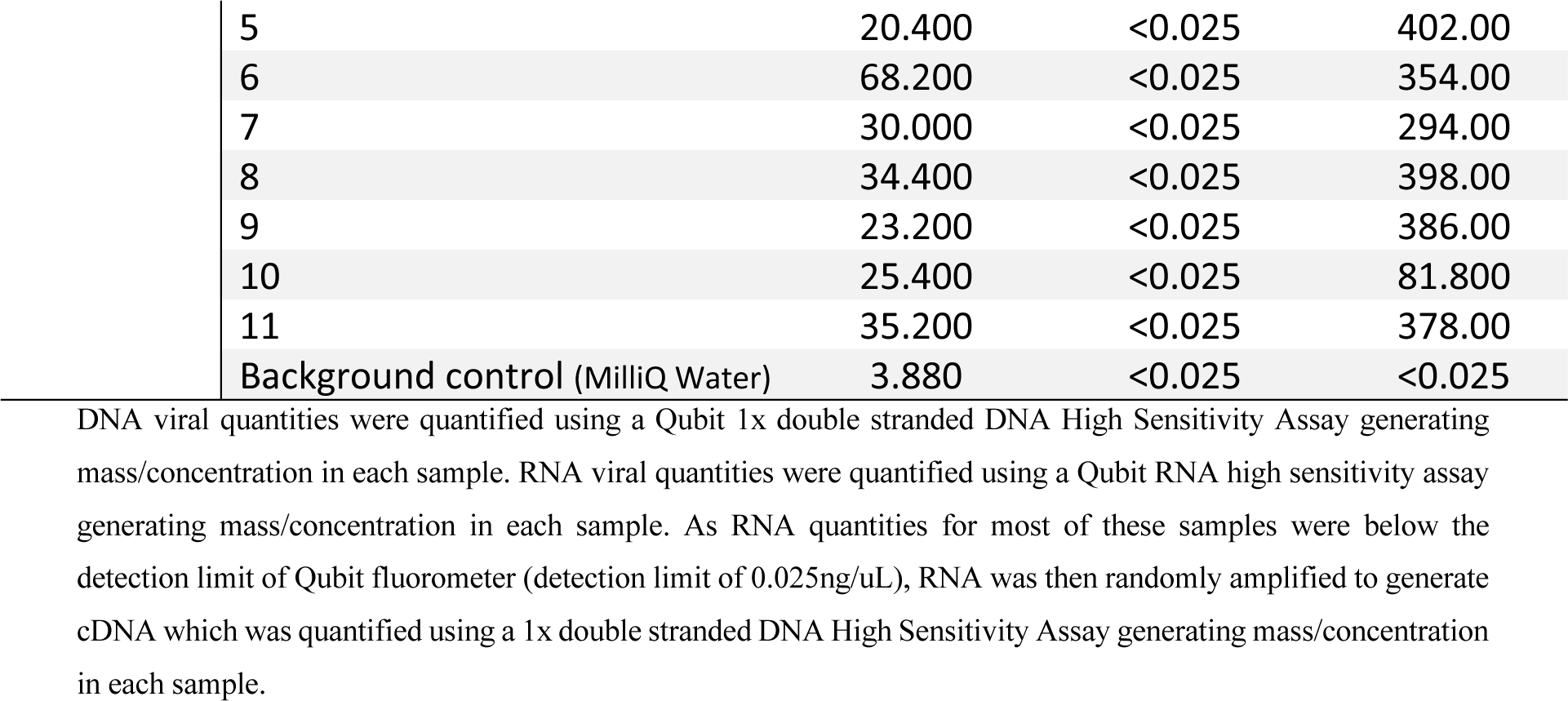
Qubit 4 Fluorometer quantification of total nucleic acids (DNA and RNA) in eleven 10L samples collected during the Spring, Summer and Fall of 2021 at different locations along the effluents of the Red and Assiniboine River within Winnipeg, MB.

| Season<br>2021 | Sample Collection Location | Viral DNA<br>(ng/uL) | Viral RNA (ng/uL) |  |
| --- | --- | --- | --- | --- |
|  |  |  | Viral RNA | cDNA |
| Spring | 1 (Control/ Undisturbed Environment) | 12.80 | <0.025 | 17.50 |
|  | 2 | 25.00 | <0.025 | 1160 |
|  | 3 | 31.00 | <0.025 | 80.00 |
|  | 4 | 6.700 | <0.025 | 143.00 |
|  | 5 | 12.60 | <0.025 | 61.80 |
|  | 6 | 18.80 | <0.025 | 25.00 |
|  | 7 | 17.10 | <0.025 | 112.0 |
|  | 8 | 58.60 | 4.00 | 47.80 |
|  | 9 | 9.660 | <0.025 | 125.00 |
|  | 10 | 13.60 | <0.025 | 84.600 |
|  | 11 | 6.580 | <0.025 | 75.600 |
|  | Background control (MilliQ Water) | 0.0208 | <0.025 | <0.025 |
| Summer | 1 (Control/ Undisturbed Environment) | 22.40 | <0.025 | 314.00 |
|  | 2 | 40.40 | 4.000 | 318.00 |
|  | 3 | 35.80 | 4.200 | 302.00 |
|  | 4 | 39.60 | 4.000 | 256.00 |
|  | 5 | 35.60 | <0.025 | 258.00 |
|  | 6 | 26.20 | <0.025 | 78.00 |
|  | 7 | 7.180 | <0.025 | 141.00 |
|  | 8 | 23.00 | <0.025 | 43.00 |
|  | 9 | 31.00 | 4.400 | 314.00 |
|  | 10 | 62.600 | 4.000 | 300.00 |
|  | 11 | 102.00 | 7.340 | 320.00 |
|  | Background control (MilliQ Water) | 0.550 | <0.025 | <0.025 |
| Fall | 1 (Control/ Undisturbed Environment) | 21.800 | <0.025 | 242.00 |
|  | 2 | 32.800 | <0.025 | 348.00 |
|  | 3 | 23.000 | <0.025 | 412.00 |
|  | 4 | 23.200 | <0.025 | 326.00 |

|  |  |  |  |
| --- | --- | --- | --- |
| 5 | 20.400 | <0.025 | 402.00 |
| 6 | 68.200 | <0.025 | 354.00 |
| 7 | 30.000 | <0.025 | 294.00 |
| 8 | 34.400 | <0.025 | 398.00 |
| 9 | 23.200 | <0.025 | 386.00 |
| 10 | 25.400 | <0.025 | 81.800 |
| 11 | 35.200 | <0.025 | 378.00 |
| Background control (MilliQ Water) | 3.880 | <0.025 | <0.025 |

DNA viral quantities were quantified using a Qubit 1x double stranded DNA High Sensitivity Assay generating mass/concentration in each sample. RNA viral quantities were quantified using a Qubit RNA high sensitivity assay generating mass/concentration in each sample. As RNA quantities for most of these samples were below the detection limit of Qubit fluorometer (detection limit of 0.025ng/uL), RNA was then randomly amplified to generate cDNA which was quantified using a 1x double stranded DNA High Sensitivity Assay generating mass/concentration in each sample.

**Table S2:**
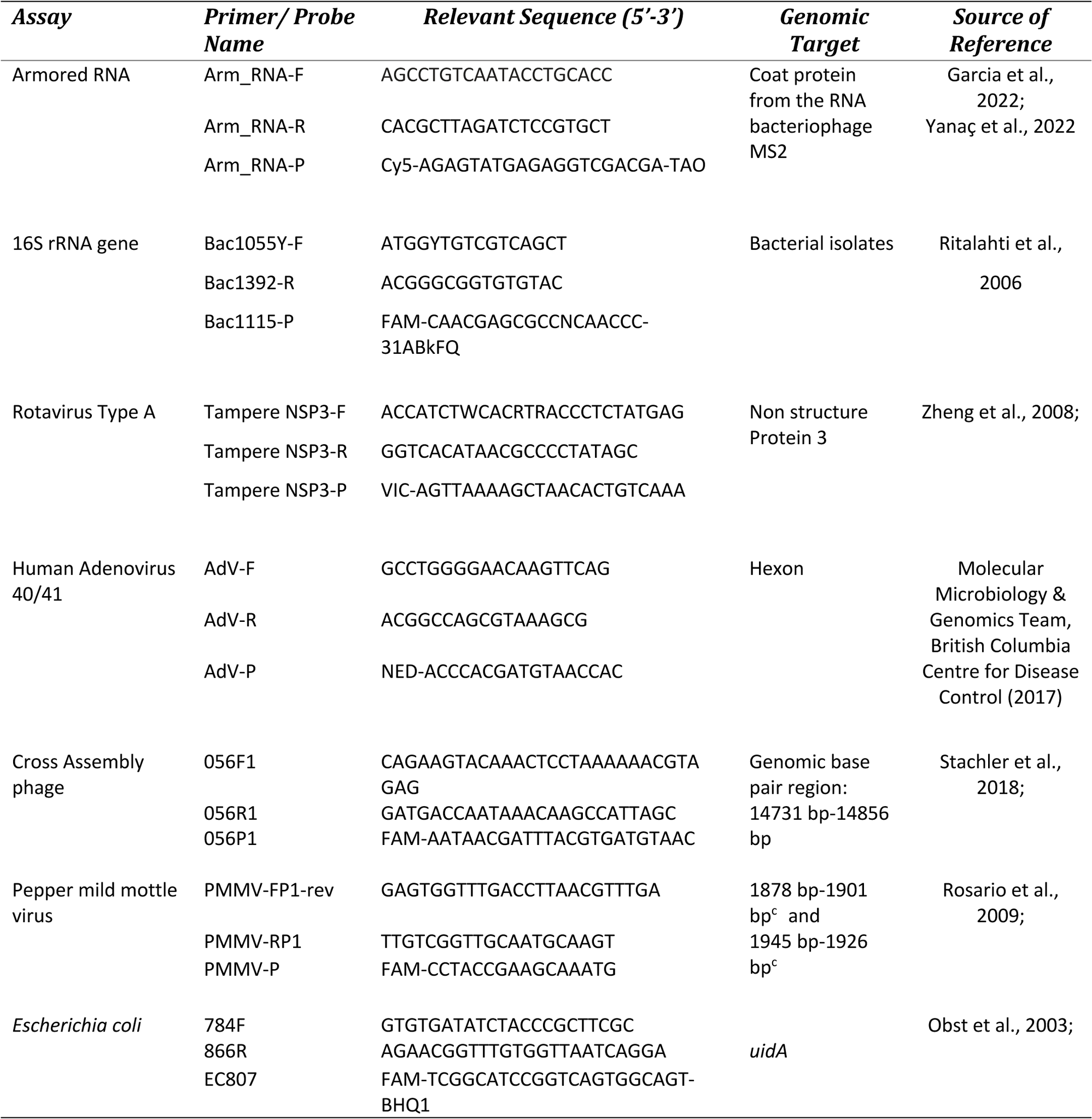
Sequences of primers and probes used in targeted q-PCR and RT–qPCR assays to screen for DNA and RNA viruses of interest.

**Table S3:**
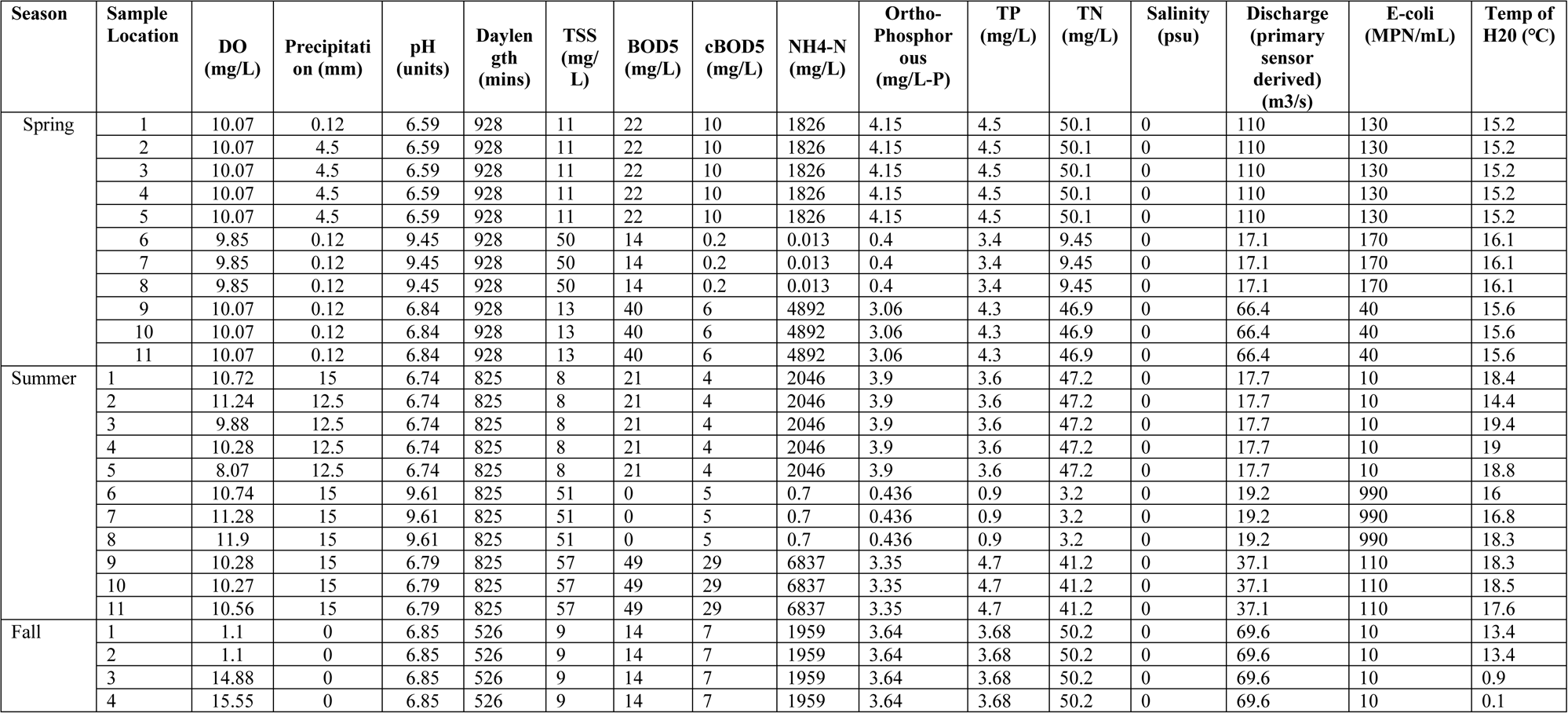
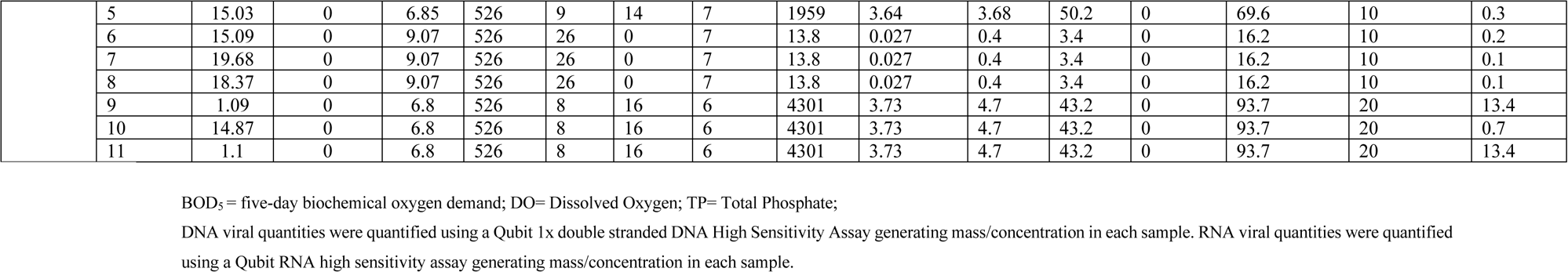
Red and Assiniboine Rivers water quality parameters present during the Spring Summer and Fall 2021.

| Season | Sample Location | DO (mg/L) | Precipitation (mm) | pH (units) | Daylength (mins) | TSS (mg/L) | BOD5 (mg/L) | cBOD5 (mg/L) | NH4-N (mg/L) | Ortho-Phosphorous (mg/L-P) | TP (mg/L) | TN (mg/L) | Salinity (psu) | Discharge (primary sensor derived) (m3/s) | E-coli (MPN/mL) | Temp of H2O (°C) |
| --- | --- | --- | --- | --- | --- | --- | --- | --- | --- | --- | --- | --- | --- | --- | --- | --- |
| Spring | 1 | 10.07 | 0.12 | 6.59 | 928 | 11 | 22 | 10 | 1826 | 4.15 | 4.5 | 50.1 | 0 | 110 | 130 | 15.2 |
|  | 2 | 10.07 | 4.5 | 6.59 | 928 | 11 | 22 | 10 | 1826 | 4.15 | 4.5 | 50.1 | 0 | 110 | 130 | 15.2 |
|  | 3 | 10.07 | 4.5 | 6.59 | 928 | 11 | 22 | 10 | 1826 | 4.15 | 4.5 | 50.1 | 0 | 110 | 130 | 15.2 |
|  | 4 | 10.07 | 4.5 | 6.59 | 928 | 11 | 22 | 10 | 1826 | 4.15 | 4.5 | 50.1 | 0 | 110 | 130 | 15.2 |
|  | 5 | 10.07 | 4.5 | 6.59 | 928 | 11 | 22 | 10 | 1826 | 4.15 | 4.5 | 50.1 | 0 | 110 | 130 | 15.2 |
|  | 6 | 9.85 | 0.12 | 9.45 | 928 | 50 | 14 | 0.2 | 0.013 | 0.4 | 3.4 | 9.45 | 0 | 17.1 | 170 | 16.1 |
|  | 7 | 9.85 | 0.12 | 9.45 | 928 | 50 | 14 | 0.2 | 0.013 | 0.4 | 3.4 | 9.45 | 0 | 17.1 | 170 | 16.1 |
|  | 8 | 9.85 | 0.12 | 9.45 | 928 | 50 | 14 | 0.2 | 0.013 | 0.4 | 3.4 | 9.45 | 0 | 17.1 | 170 | 16.1 |
|  | 9 | 10.07 | 0.12 | 6.84 | 928 | 13 | 40 | 6 | 4892 | 3.06 | 4.3 | 46.9 | 0 | 66.4 | 40 | 15.6 |
|  | 10 | 10.07 | 0.12 | 6.84 | 928 | 13 | 40 | 6 | 4892 | 3.06 | 4.3 | 46.9 | 0 | 66.4 | 40 | 15.6 |
|  | 11 | 10.07 | 0.12 | 6.84 | 928 | 13 | 40 | 6 | 4892 | 3.06 | 4.3 | 46.9 | 0 | 66.4 | 40 | 15.6 |
| Summer | 1 | 10.72 | 15 | 6.74 | 825 | 8 | 21 | 4 | 2046 | 3.9 | 3.6 | 47.2 | 0 | 17.7 | 10 | 18.4 |
|  | 2 | 11.24 | 12.5 | 6.74 | 825 | 8 | 21 | 4 | 2046 | 3.9 | 3.6 | 47.2 | 0 | 17.7 | 10 | 14.4 |
|  | 3 | 9.88 | 12.5 | 6.74 | 825 | 8 | 21 | 4 | 2046 | 3.9 | 3.6 | 47.2 | 0 | 17.7 | 10 | 19.4 |
|  | 4 | 10.28 | 12.5 | 6.74 | 825 | 8 | 21 | 4 | 2046 | 3.9 | 3.6 | 47.2 | 0 | 17.7 | 10 | 19 |
|  | 5 | 8.07 | 12.5 | 6.74 | 825 | 8 | 21 | 4 | 2046 | 3.9 | 3.6 | 47.2 | 0 | 17.7 | 10 | 18.8 |
|  | 6 | 10.74 | 15 | 9.61 | 825 | 51 | 0 | 5 | 0.7 | 0.436 | 0.9 | 3.2 | 0 | 19.2 | 990 | 16 |
|  | 7 | 11.28 | 15 | 9.61 | 825 | 51 | 0 | 5 | 0.7 | 0.436 | 0.9 | 3.2 | 0 | 19.2 | 990 | 16.8 |
|  | 8 | 11.9 | 15 | 9.61 | 825 | 51 | 0 | 5 | 0.7 | 0.436 | 0.9 | 3.2 | 0 | 19.2 | 990 | 18.3 |
|  | 9 | 10.28 | 15 | 6.79 | 825 | 57 | 49 | 29 | 6837 | 3.35 | 4.7 | 41.2 | 0 | 37.1 | 110 | 18.3 |
|  | 10 | 10.27 | 15 | 6.79 | 825 | 57 | 49 | 29 | 6837 | 3.35 | 4.7 | 41.2 | 0 | 37.1 | 110 | 18.5 |
|  | 11 | 10.56 | 15 | 6.79 | 825 | 57 | 49 | 29 | 6837 | 3.35 | 4.7 | 41.2 | 0 | 37.1 | 110 | 17.6 |
| Fall | 1 | 1.1 | 0 | 6.85 | 526 | 9 | 14 | 7 | 1959 | 3.64 | 3.68 | 50.2 | 0 | 69.6 | 10 | 13.4 |
|  | 2 | 1.1 | 0 | 6.85 | 526 | 9 | 14 | 7 | 1959 | 3.64 | 3.68 | 50.2 | 0 | 69.6 | 10 | 13.4 |
|  | 3 | 14.88 | 0 | 6.85 | 526 | 9 | 14 | 7 | 1959 | 3.64 | 3.68 | 50.2 | 0 | 69.6 | 10 | 0.9 |
|  | 4 | 15.55 | 0 | 6.85 | 526 | 9 | 14 | 7 | 1959 | 3.64 | 3.68 | 50.2 | 0 | 69.6 | 10 | 0.1 |
|  | 5 | 15.03 | 0 | 6.85 | 526 | 9 | 14 | 7 | 1959 | 3.64 | 3.68 | 50.2 | 0 | 69.6 | 10 | 0.3 |
|  | 6 | 15.09 | 0 | 9.07 | 526 | 26 | 0 | 7 | 13.8 | 0.027 | 0.4 | 3.4 | 0 | 16.2 | 10 | 0.2 |
|  | 7 | 19.68 | 0 | 9.07 | 526 | 26 | 0 | 7 | 13.8 | 0.027 | 0.4 | 3.4 | 0 | 16.2 | 10 | 0.1 |
|  | 8 | 18.37 | 0 | 9.07 | 526 | 26 | 0 | 7 | 13.8 | 0.027 | 0.4 | 3.4 | 0 | 16.2 | 10 | 0.1 |
|  | 9 | 1.09 | 0 | 6.8 | 526 | 8 | 16 | 6 | 4301 | 3.73 | 4.7 | 43.2 | 0 | 93.7 | 20 | 13.4 |
|  | 10 | 14.87 | 0 | 6.8 | 526 | 8 | 16 | 6 | 4301 | 3.73 | 4.7 | 43.2 | 0 | 93.7 | 20 | 0.7 |
|  | 11 | 1.1 | 0 | 6.8 | 526 | 8 | 16 | 6 | 4301 | 3.73 | 4.7 | 43.2 | 0 | 93.7 | 20 | 13.4 |

BOD<sub>5</sub> = five-day biochemical oxygen demand; DO= Dissolved Oxygen; TP= Total Phosphate;
DNA viral quantities were quantified using a Qubit 1x double stranded DNA High Sensitivity Assay generating mass/concentration in each sample. RNA viral quantities were quantified
using a Qubit RNA high sensitivity assay generating mass/concentration in each sample.

**FIG S1:**
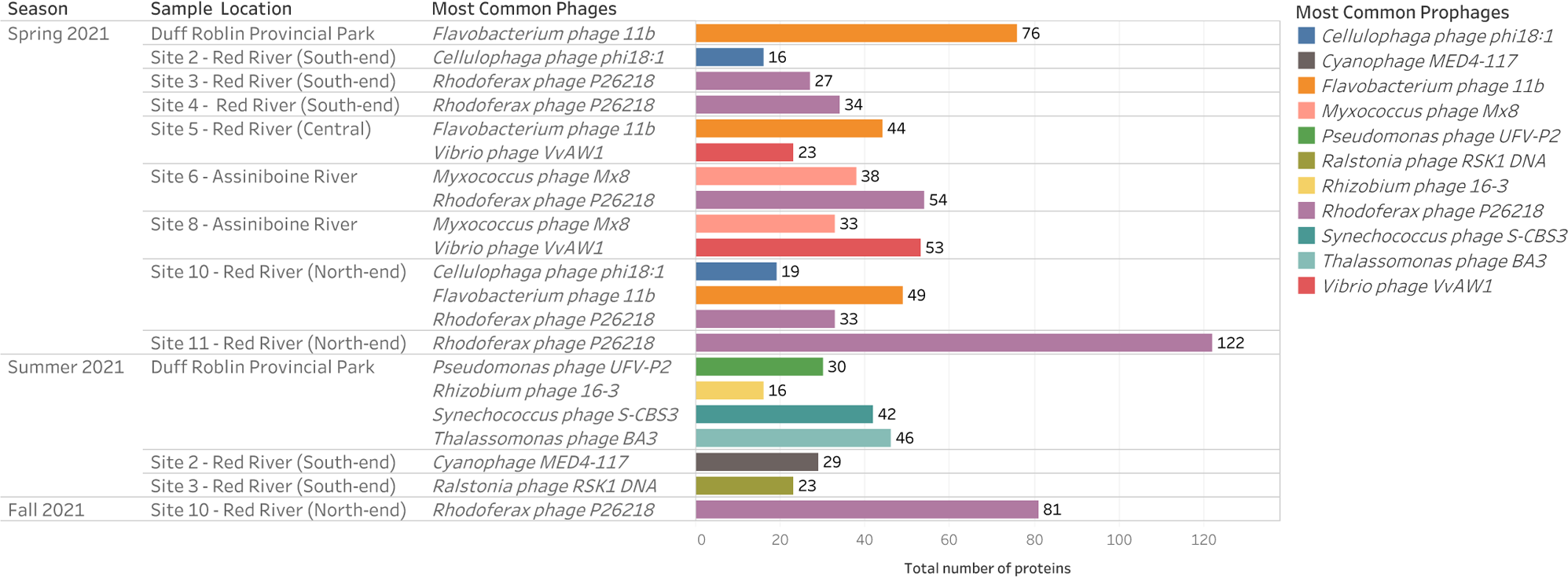
Comparison of the 2021 DNA and RNA phage composition during the Spring, Summer and Fall of 2021 that were identified from PHASTER with complete scores and intact regions

**FIG S2:**
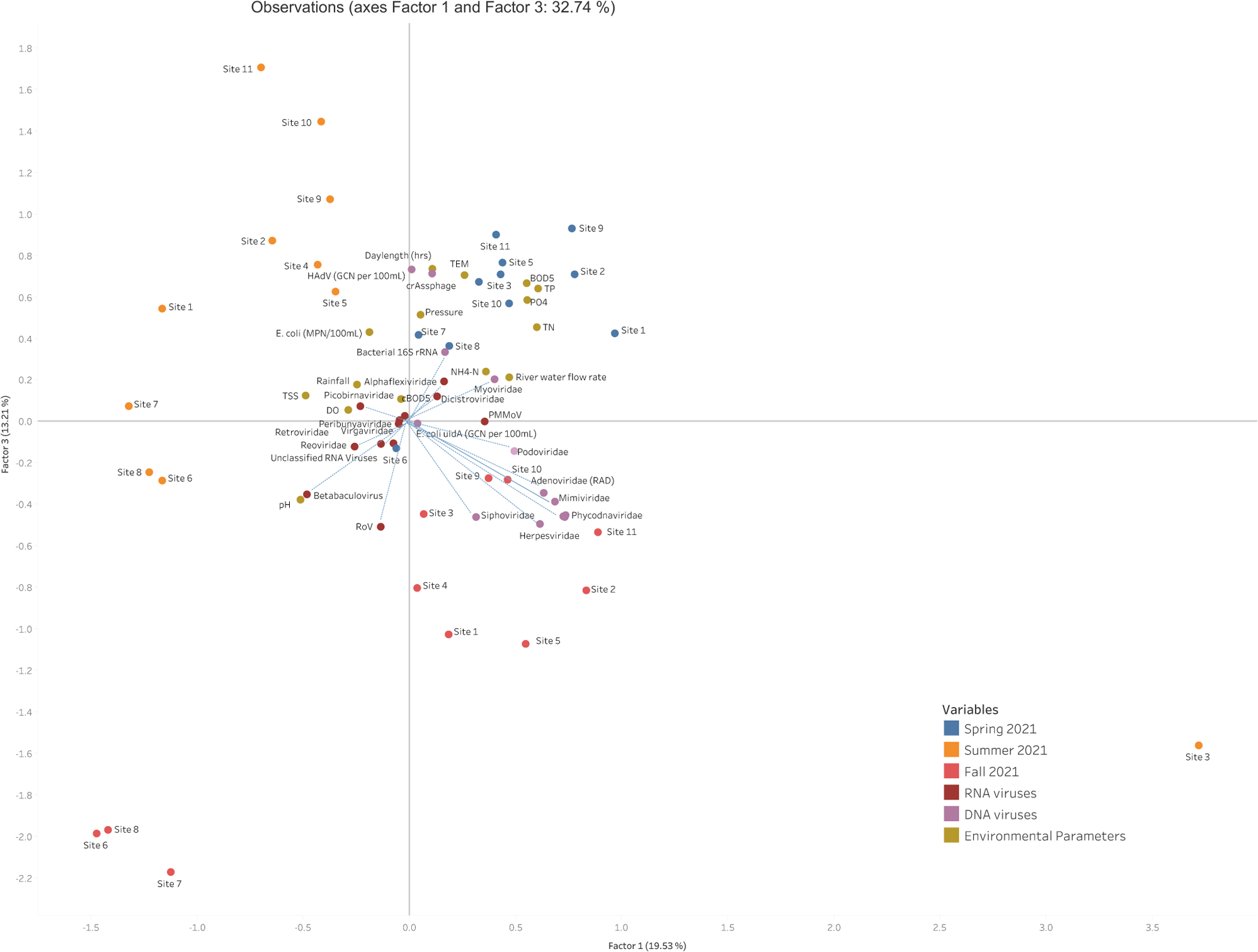
Factor analysis of viral DNA and RNA families and environmental variables observed at each sample collection site (1-11) during the Spring (blue dots), Summer (orange dots) and Fall (red dots) of 2021. Factor 1 represents land use which incorporates urban influenced watersheds on DNA viruses and agricultural influenced watershed on RNA viruses while factor 3 represents the environmental water quality parameter flow rate. TSS, total suspension solids; DO, dissolved oxygen; TEM, temperature; cBOD_5_, carbonaceous biochemical oxygen demand; BOD, biochemical oxygen demand; TP, total phosphorous; TN, total nitrogen; NH4-N, ammonia, PO_4_, phosphorous; *E. coli* (MPN/100mL), *E.coli* colony forming unit counts. Blue dashed lines represent factor loading values for viral families.

**FIG S3:**
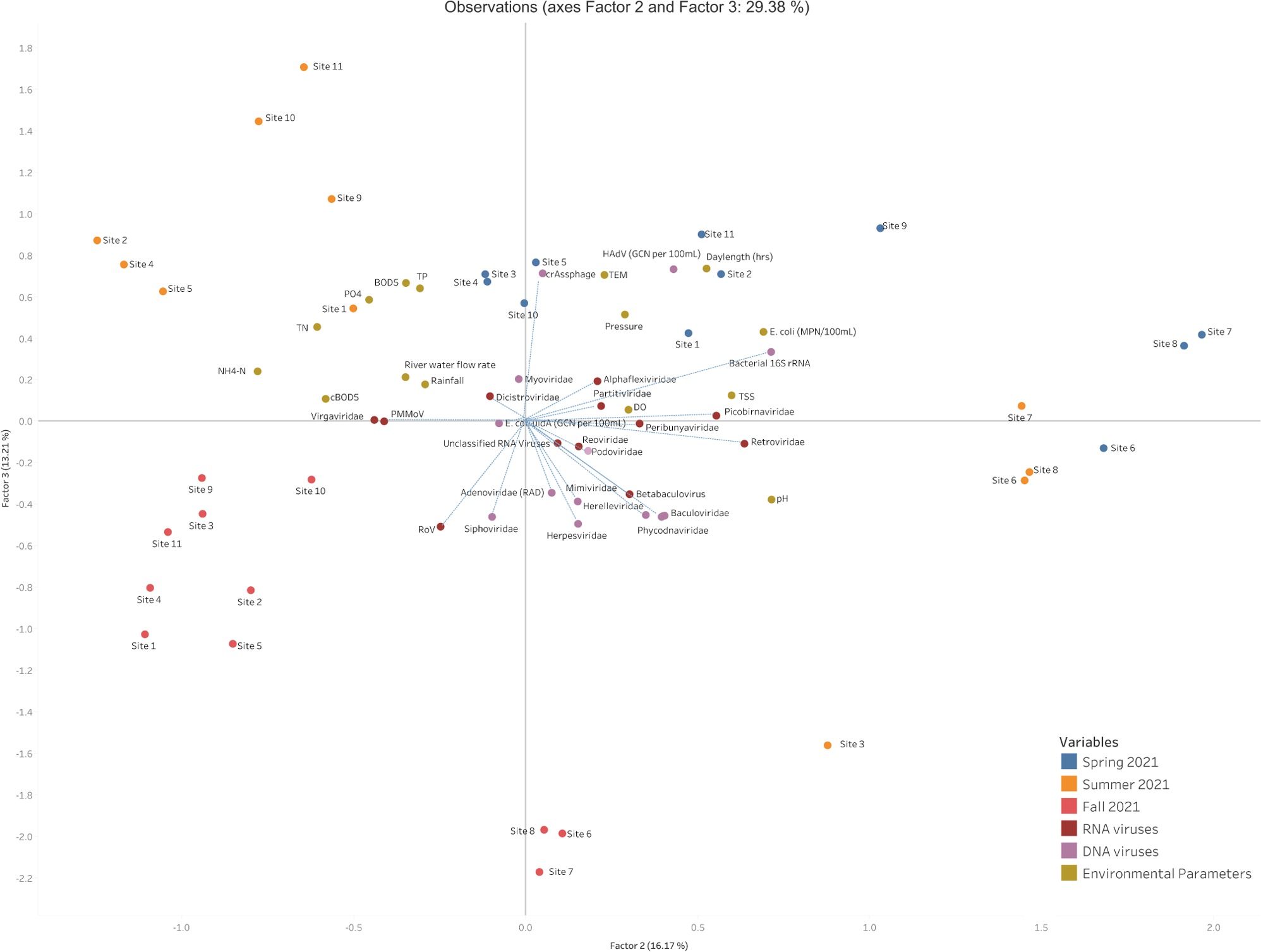
Factor analysis of viral DNA and RNA families and environmental variables observed at each sample collection site (1-11) during the Spring (blue dots), Summer (orange dots) and Fall (red dots) of 2021. Factor 2 represents the water quality parameters daylength and temperature while factor 3 represents the environmental water quality parameter flow rate. TSS, total suspension solids; DO, dissolved oxygen; TEM, temperature; cBOD_5_, carbonaceous biochemical oxygen demand; BOD, biochemical oxygen demand; TP, total phosphorous; TN, total nitrogen; NH4-N, ammonia, PO_4_, phosphorous; *E. coli* (MPN/100mL), *E.coli* colony forming unit counts. Blue dashed lines represent factor loading values for viral families.

**FIG. S4:**
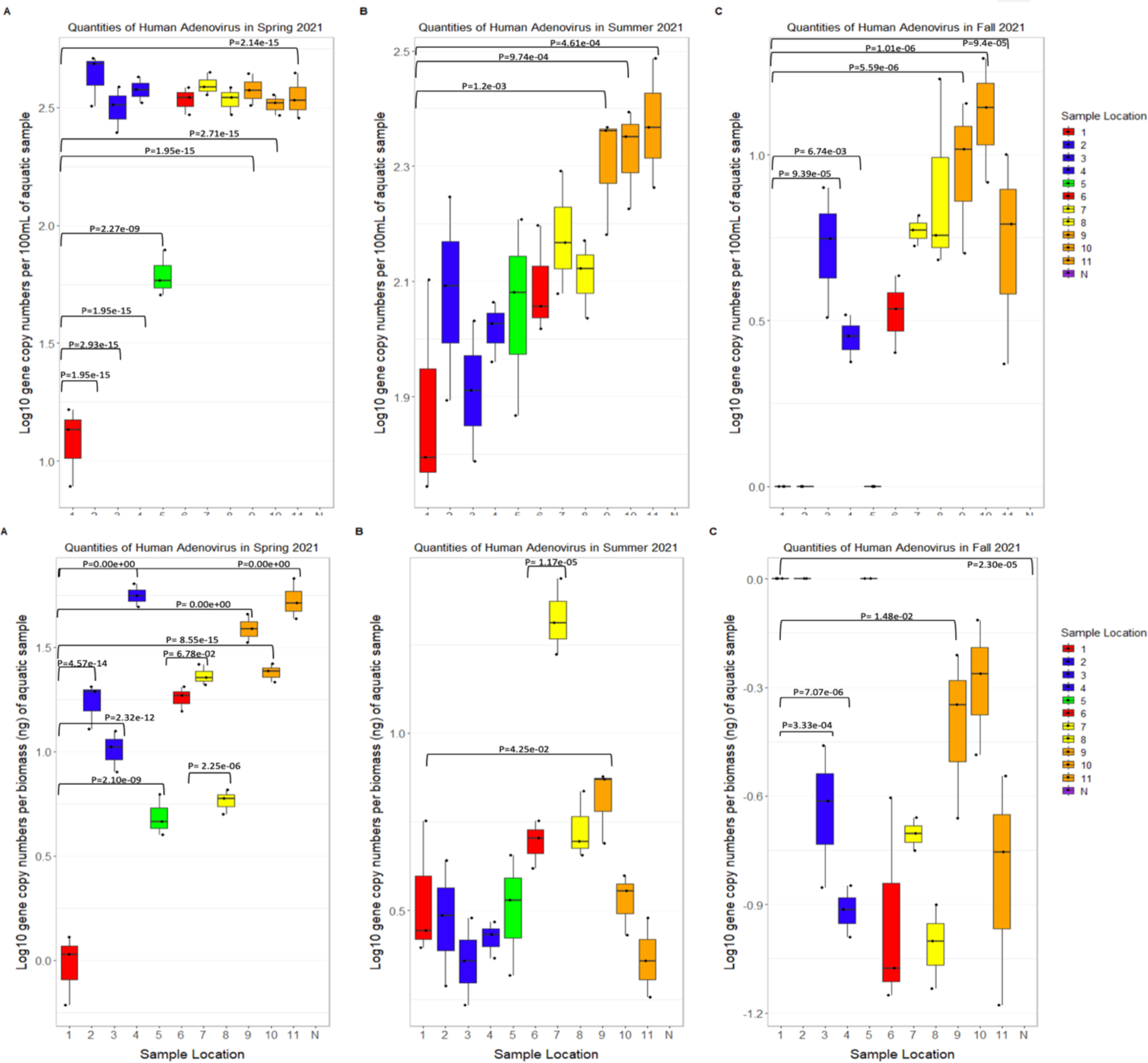
Abundance of *Human Adenovirus* per volume (100mL) (top panel) and per biomass (ng) (bottom panel) in aquatic samples 1–11 collected from the Red and Assiniboine rivers of Winnipeg, MB during (A) Spring, (B) Summer and (C) Fall 2021.

**FIG. S5:**
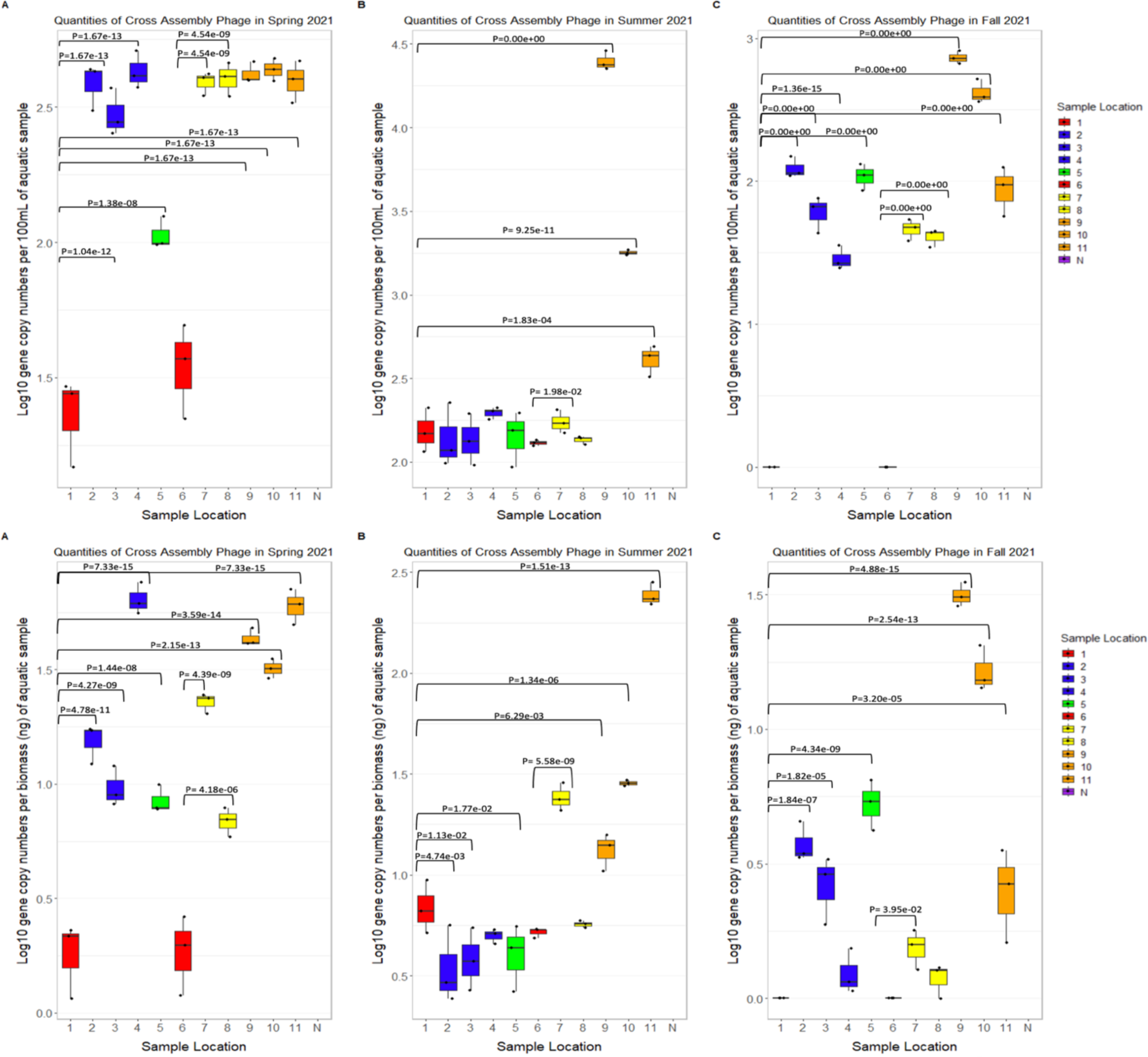
Abundance of *Cross-Assembly Phage* per volume (100mL) (top panel) and per biomass (ng) (bottom panel) in aquatic samples 1–11 collected from the Red and Assiniboine rivers of Winnipeg, MB during (A) Spring, (B) Summer and (C) Fall 2021.

**FIG S6:**
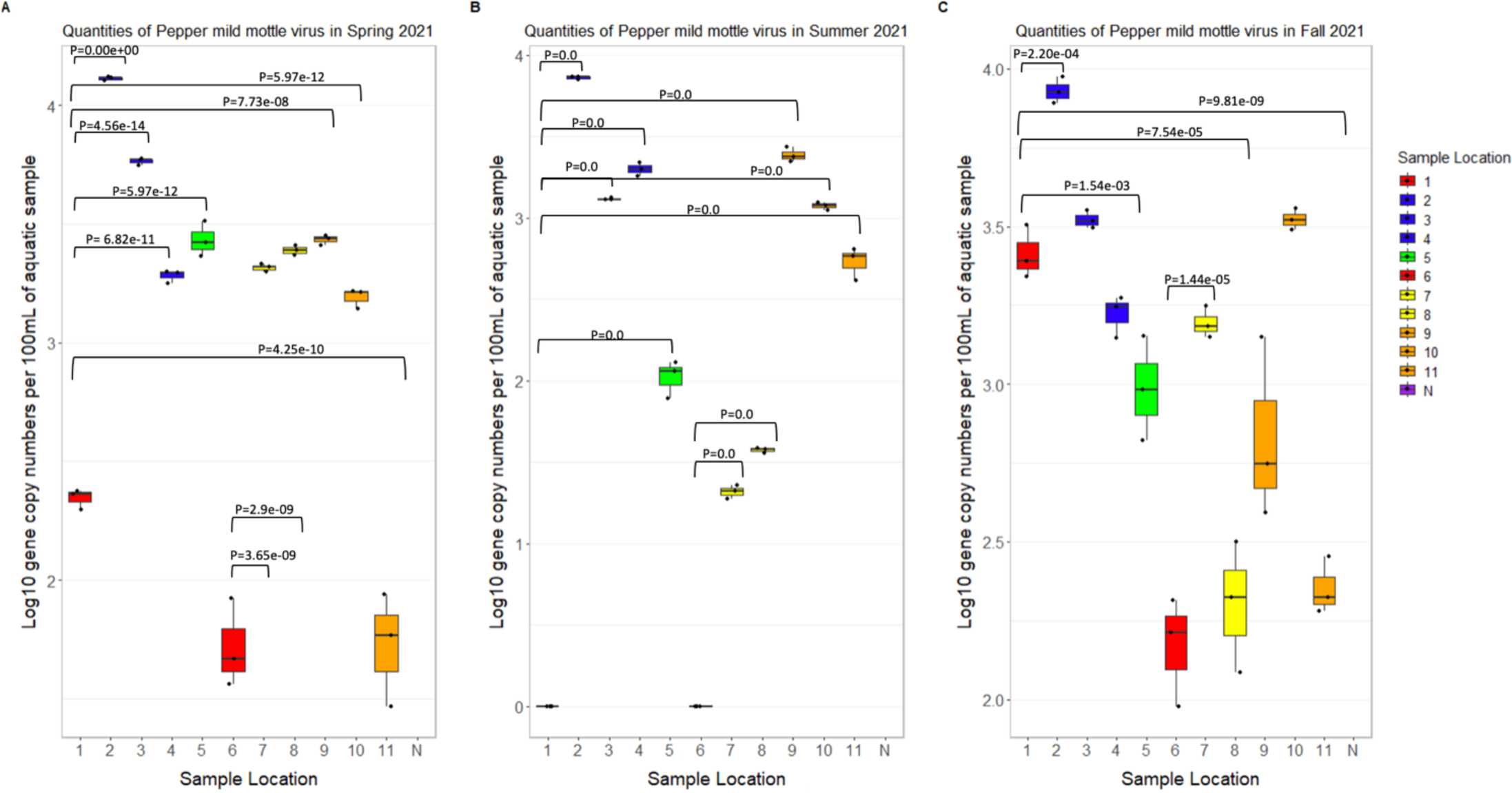
Abundance of *Pepper-mild mottle virus* per volume (100 mL) in aquatic samples 1–11 collected from the Red and Assiniboine rivers of Winnipeg, MB during (A) Spring, (B) Summer and (C) Fall 2021.

**FIG S7:**
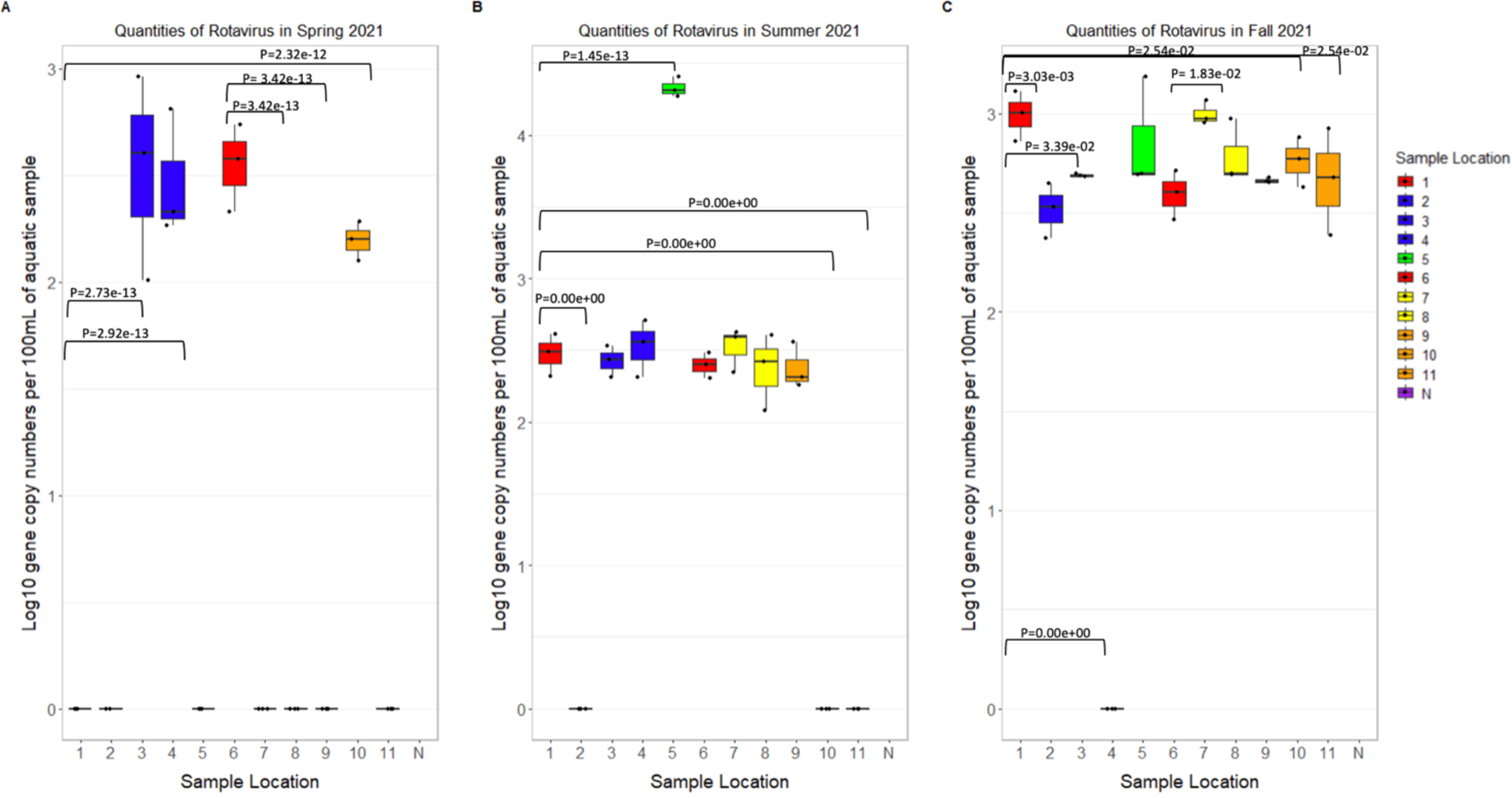
Abundance of *Rotavirus* per volume (100 mL) in aquatic samples 1–11 collected from the Red and Assiniboine rivers of Winnipeg, MB during (A) Spring, (B) Summer and (C) Fall 2021.

